# Minimal Perturbation of Activation Loop Dynamics Rewires Kinase Signaling

**DOI:** 10.1101/2025.10.15.682502

**Authors:** Prashant Jain, Dariia Yehorova, Ririn Rahamala Febri, Ben E. Clifton, Andrei Demkiv, Genichiro Uechi, Michael Robinson, Shina Caroline Lynn Kamerlin, Mariko Okada, Akira Imamoto, Paola Laurino

**Affiliations:** Protein Engineering and Evolution Unit, Okinawa Institute of Science and Technology, Okinawa, Japan; School of Chemistry and Biochemistry and of Chemical and Biological Engineering, Georgia Institute of Technology, Atlanta, GA, USA; Institute for Protein Research, Osaka University, Suita, Japan; Department of Cell and Molecular Biology, Uppsala University, Sweden; School of Chemical and Biological Engineering, Georgia Institute of Technology, Atlanta, GA; Department of Chemistry, Lund University, Lund, Sweden; The Ben May Department, The University of Chicago, Chicago, IL, USA

## Abstract

Enzymes are central to life, with their catalytic activity often shaped by the dynamic conformations of regulatory loops. In hub enzymes such as tyrosine kinases, the activation loop critically controls substrate specificity, catalytic efficiency, and downstream signaling, shaping cellular fate. Yet, the molecular mechanisms by which loop dynamics encode these functions remain incompletely understood. Here, we used SRC kinase as a model to dissect how minimal perturbations of the activation loop reprogram kinase behavior. By generating and characterizing multiple variants, we identified a triple-deletion mutant with altered loop dynamics. Structural and biochemical analyses revealed that this variant explores distinct loop conformations and exhibits a subtle shift in substrate preference toward more acidic motifs. These fine-tuned conformational changes translated into specific cellular signaling outcomes, as demonstrated by phosphoproteomic profiling. Comparative analysis across species further showed that nature exploits similar loop remodeling strategies to modulate kinase function. Together, our findings provide a blueprint for rationally tuning kinase activity and offer a generalizable framework for rewiring signaling pathways in diverse cellular contexts.

## Introduction

Controlling cellular signaling pathways holds immense potential for both reengineering biological function^1,2^ and elucidating the mechanisms underlying physiological impacts^3,4^. Central to these networks are enzymes^5^, particularly kinases, which catalyze reactions that regulate the functional activity^6^, localization^7^, and interactions of signaling molecules^8^. While signal transduction mechanisms such as phosphorylation^9^, dephosphorylation^10^, proteolysis^11^, ubiquitination^12^, and G-protein-coupled receptor (GPCR) signaling^13^, have been extensively studied, many molecular aspects of kinase regulation remain unsolved, limiting our ability to precisely manipulate signaling.

Protein engineering offers powerful strategies to rewire signaling nodes^14,15^, yet current approaches such as orthogonal domains^16^, rewired phosphorylation circuits^17^, chemically inducible systems^18^, and optogenetic systems^19^, often rely on substantial modifications of the protein backbone or non-native components, which can disrupt endogenous network architecture^20^. In contrast, understanding how nascent kinase domains achieve substrate specificity^21,22^ and signaling fidelity provides a minimally invasive route to modulate signaling at its source.

The activation loop (A-loop) is a dynamic and disordered structural element within the kinase catalytic core^23^ that critically governs substrate recognition^24^, autophosphorylation^25^, and conformational transitions between inactive and active states. In SRC, a ubiquitous non-receptor tyrosine kinase and hub of multiple signaling pathways^26^, the A-loop undergoes autophosphorylation at Y416^27^, triggering structural rearrangements that unlock catalytic activity^28^ (Figures 1a and 1b). Despite extensive characterization of its structural role^29,30^, and conformational dynamics^31,32^, the A-loop remains largely untapped as a target for precise, minimal-perturbation engineering to fine-tune kinase activity and cellular signaling.

**Figure 1:**
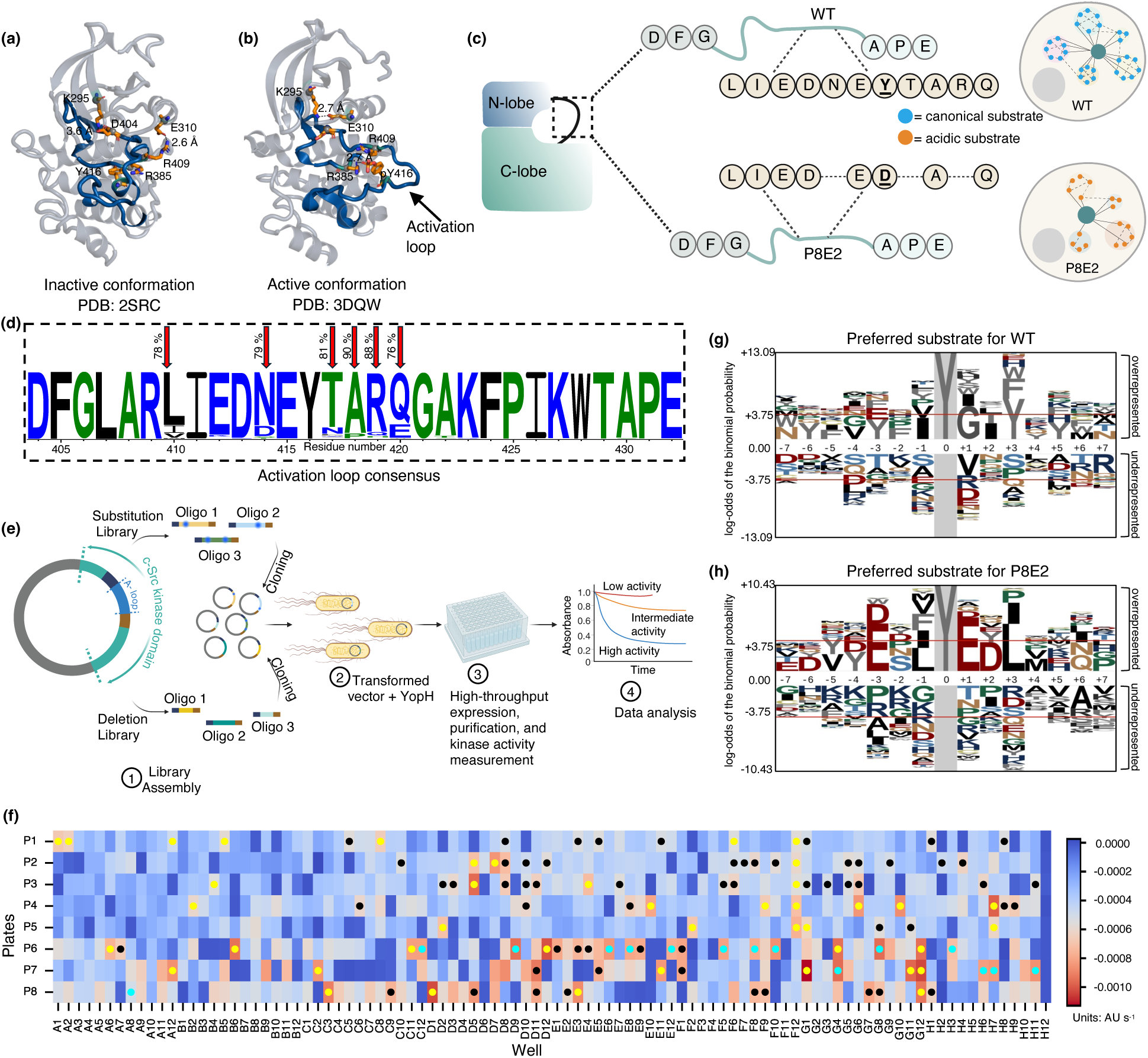
Activation Loop Remodeling Rewires SRC Kinase Function and Substrate Specificity. **(a and b)** Structural comparison of the SRC kinase domain in its inactive conformation (**a**, PDB: 2SRC^87^) and active conformation (**b**, PDB: 3DQW^37^), highlighting conformational changes in the activation loop (A-loop) shown in deep blue. Key residues involved in hydrogen bonding are depicted as orange spheres and sticks, with hydrogen bonds represented as dotted lines. **(c)** Shortening of the activation loop in SRC kinase alters cellular signaling and reduces substrate specificity. In the P8E2 variant, the canonical autophosphorylation site Y416 (chicken SRC numbering) is substituted with Asp (bold and underlined), and three residues within the loop are deleted (shown as dashes). **(d)** Sequence logo of activation loop deduced from MSA created with SRC kinase and related sequences (496 sequences, Supplementary Data 1). Red arrows indicate the positions selected for library construction, with labels showing the percentage sequence identity at each site. **(e)** Schematic of library assembly and screening. Substitution and deletion oligonucleotides (Supplementary Data 2) were cloned separately into the pET28a (+) vector and co-transformed with YopH in the pCDFDuet-1 vector into BL21 *E. coli* by electroporation. Approximately 800 random colonies were picked and expressed in 96-deep-well plates for high-throughput screening. Purified lysates were assayed for kinase activity at a fixed ATP concentration to compare variant activities. **(f)** Kinase activities were measured using a coupled enzymatic assay involving pyruvate kinase and lactate dehydrogenase. In this assay, oxidation of NADH results in a decrease in absorbance at 340 nm, and the rate of this decrease is directly proportional to kinase activity (a faster decline corresponds to higher activity). Heatmap shows the rate of NADH consumption within the linear range (250– 300 s) for the substitution library (P1 to P5) and deletion library (P6 to P8). Cells highlighted with yellow dots represent WT clones randomly picked from each plate. Cyan dots mark duplicate colonies randomly picked, and cells marked with black dots denote mutants that were pooled, re-expressed, and re-analyzed together in a new 96-deep-well plate. **(g and h)** Phospho-pLogos displaying preferred substrates identified from in vitro library screening for the WT **(g)** and P8E2 mutant **(h)**. To generate these logos, peptides with average enrichment scores >1.5 (strong substrates) across three biological replicates were first selected. For WT, peptides that also had enrichment scores >1.5 for P8E2 were removed, leaving 88 peptides uniquely enriched for WT. For P8E2, peptides that also had enrichment scores >1.5 for WT were removed, leaving 75 peptides uniquely enriched for P8E2. These exclusive sets, derived from the total library of 2,587 peptides, reveal distinct substrate preferences between WT and P8E2. Logos generated without this comparative filtering, showing all peptides with enrichment scores >1.5 for each variant independently, are provided in Extended Figure 2.

Here, we systematically analyzed SRC and related kinase sequences to identify residues in the A-loop amenable to engineering. We generated ~190 variants, revealing a spectrum of effects on activity, from inactivation to subtle functional shifts. From this library, the P8E2 variant, harboring three deletions (ΔN414, ΔT417, ΔR419) and a substitution (Y416D), exhibited a modest increase in catalytic efficiency and a measurable shift in substrate preference. Structural and molecular dynamics analyses showed that P8E2 explores distinct loop conformations, correlating with altered cellular signaling outcomes in HEK293T cells, as confirmed by phosphoproteomic and expression profiling. Phylogenetic analysis further suggests that nature exploits similar loop remodeling strategies to fine-tune kinase function across species. Together, these findings demonstrate that minimal perturbations of the activation loop provide a generalizable framework for rewiring kinase activity and controlling signaling with precision (Figure 1c).

## Results

### Activation Loop Engineering Reveals How Strategic Deletions Enhance Catalytic Efficiency and Bypass Autophosphorylation

To test the role of autophosphorylation in regulating kinase activity, we introduced the Y416D mutation in SRC. Acidic substitutions such as Asp or Glu are widely used as phosphorylation mimics for serine, threonine, and tyrosine residues because their negative charge can substitute for that of a phosphate group^33,34^. We hypothesized that replacing Y416 with Asp would constrain activation loop dynamics by bypassing the need for phosphorylation and eliminating the requirement to toggle between phosphorylated (open) and unphosphorylated (closed) states. Such a constraint was expected to simplify regulatory control and yield a simplified signaling response (Supplementary Note 1). However, kinetic analysis revealed that the Y416D variant exhibited an approximately two-fold reduction in catalytic efficiency (3400 M^−1^ s^−1^), compared to wild-type (WT) SRC (7900 M^−1^ s^−1^) (Table 1, Extended Figure 1a), indicating that acidic substitutions cannot recapitulate the stabilizing effects of phosphorylation.

**Table 1:**
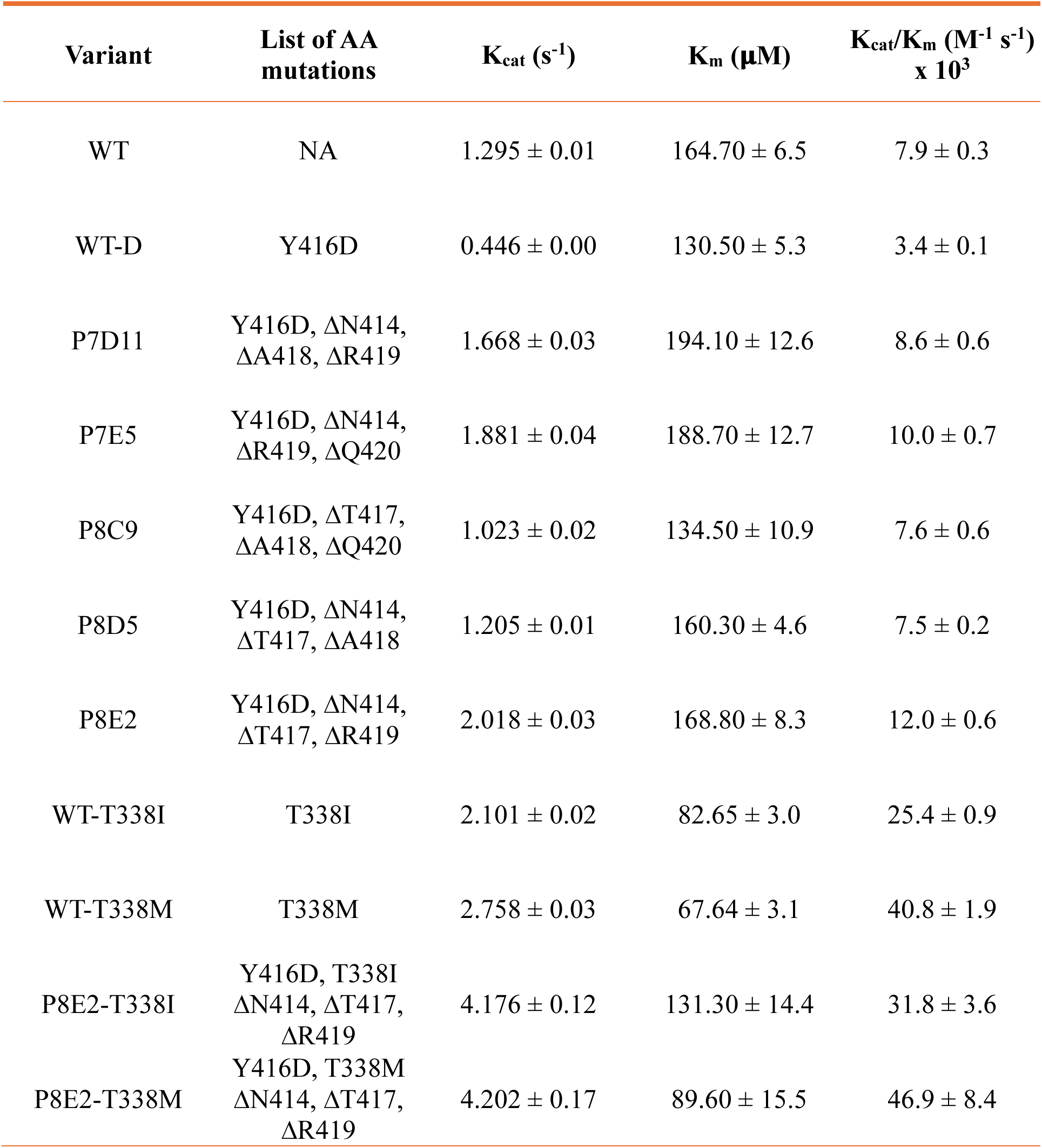
Kinetic parameters of SRC variants using ATP as the substrate. Parameters were obtained from initial rate measurements with a synthetic peptide in a coupled enzymatic assay. Values represent best-fit estimates ± error derived from the 95% confidence interval from four independent measurements

Building on this observation, we engineered a library of activation loop mutants to identify variants that could bypass autophosphorylation while preserving catalytic efficiency. A multiple sequence alignment (MSA) of SRC homologs, SRC family kinases, and related kinases (~500 sequences) revealed that while the activation loop is generally highly conserved (sequence identity 97%), a subset of six positions within this region displays increased variability, with an sequence identity of 82% (Figure 1d). Based on this, we created a library of single and double mutants, targeting the less conserved residues in the activation loop (six residues, L410, N414, T417, A418, R419 and Q420, along with mutants featuring deletions of A-loop residues in all possible combinations (Table S1).

Following cloning, high-throughput expression, purification, and kinetic analysis (Figure 1e), we identified 60 mutants with catalytic efficiencies ranging from 65% to 97% of WT SRC, comparable to the activity of WT enzyme (Figure 1f). The top five candidates, all containing three deletions in combination with Y416D, were selected for large-scale expression. Michaelis–Menten parameters for these variants are presented in Table 1. The most promising variant, P8E2, showed deletions at N414, T417 and R419 positions in the loop, exhibiting catalytic efficiency very similar to WT SRC (12000 M^−1^s^−1^ for P8E2 vs 7900 M^−1^s^−1^ for WT) while bypassing the autophosphorylation (Table 1 and Extended Figure 1a).

In addition, we examined the role of the conserved gatekeeper residue, T338, which sits in the ATP-binding pocket and is frequently mutated in cancers^35^. Substitutions at this position (e.g., T338M and T338I) are known to enhance kinase activity by destabilizing the inactive state^36^ and favoring active conformations^37^. Based on this, we hypothesized that introducing gatekeeper mutations could expand the conformational space of SRC and thereby influence signaling dynamics. Consistent with this expectation, biochemical assays revealed 3- to 5-fold increase in catalytic efficiency of the gatekeeper variants, driven by improvements in both turnover number and substrate affinity (Table 1, Extended Figure 1b and 1c). These findings suggest that gatekeeper mutations shift the kinase towards conformations capable of efficient substrate phosphorylation. Further mechanistic details are provided in Supplementary Note 2.

### A Subtle Yet Significant Shift in the Substrate Specificities Between WT And P8E2

To investigate whether the introduced mutations altered substrate preference, we performed a high-throughput specificity screen using a bacterial surface-display library containing 15-residue peptides with known human tyrosine phosphorylation sites (“Human-pTyr” library), following the general strategy established by Kuriyan et al^38,39^. The library was phosphorylated by the kinase of interest, labeled with a fluorescent pan-phosphotyrosine antibody, sorted by FACS, and analyzed via deep sequencing.

To visualize substrate preference and position-dependent amino acid selection for WT, P8E2, and their corresponding gatekeeper mutants, we generated sequence probability logos (phospho-pLogos)^40^ based on the peptide enrichment scores. The overall phospho-pLogos were similar across variants, with minor differences such as glycine enrichment at the P+1 position in WT and a slight preference for aspartate in P8E2 (Extended Figure 2a and 2b).

To detect more subtle shifts in specificity, we divided the peptides into two groups: one with strong WT substrate (enrichment score > 1.5) and weak P8E2 substrate (enrichment score < 1.5), and the other with the opposite specificity. Phospho-pLogos generated for these groups revealed distinct differences between the variants. For WT, there was an overrepresentation of aromatic residues at both the C- and N-terminal positions relative to the phosphorylation site, particularly at P+3 and P-3 (Figure 1g). In contrast, P8E2 exhibited a significant overrepresentation of negatively charged residues at both C- and N-terminal positions, along with an underrepresentation of positively charged residues C-terminal to the phosphotyrosine (Figure 1h). Additionally, the P+3 position, which strongly favored aromatic residues in WT, was switched to a preference for hydrophobic residues in P8E2.

Thus, strategic deletions within the non-conserved regions of the activation loop produced measurable shifts in kinase function, demonstrating that even minimal modifications can tune the molecular interactions underlying substrate recognition and catalysis. These changes were sufficient to reprogram activity without broadly disrupting the enzyme’s overall structure or function. Interestingly, introducing the gatekeeper mutations did not further alter substrate specificity, as its phospho-pLogo closely mirrored that of the parent deletion variant, highlighting the dominant role of the activation loop in shaping kinase selectivity (Extended Figure 2c-f).

### Computational Modeling Reveals How Activation Loop Changes Control Kinase Activity and Specificity

Molecular dynamics simulations revealed the structural basis for the catalytic impairment of the Y416D SRC mutant and its restoration in the P8E2 variant. Root mean square fluctuation (RMSF) analysis of the A-loop C_α_-atoms showed that the Y416D substitution introduces pronounced A-loop flexibility compared to the WT enzyme, whereas in WT pY416 and the P8E2 variant, the A-loop appears to be less flexible (Figure 2a). In WT pY416, the loop is stabilized by a persistent salt bridge between the pY416 and R385 side chains (Figure 2b). The Y416D substitution does introduce a negative charge at this position. However, the shorter Asp side chain, as well as the −1 charge of D416 compared to the −2 charge of pY416, weakens the salt bridge to R385. This results in the loss of stabilized contact within 2-3Å and instead increases A-loop conformational disorder through the formation of transient interactions between D416 and with neighboring residues (D413, R419). In contrast, the triple deletion in P8E2 shortens the A-loop, physically positioning D416 within interaction range of R385, narrowing the preferential distance to 1.5-2.5Å and thus re-establishing the salt bridge which restores loop rigidity without the need for phosphorylation (Figure 2b). These results demonstrate that minimal perturbations in loop length can compensate for the loss of canonical regulatory contacts, establishing a link between structural rearrangements and observed changes in enzymatic activity.

**Figure 2:**
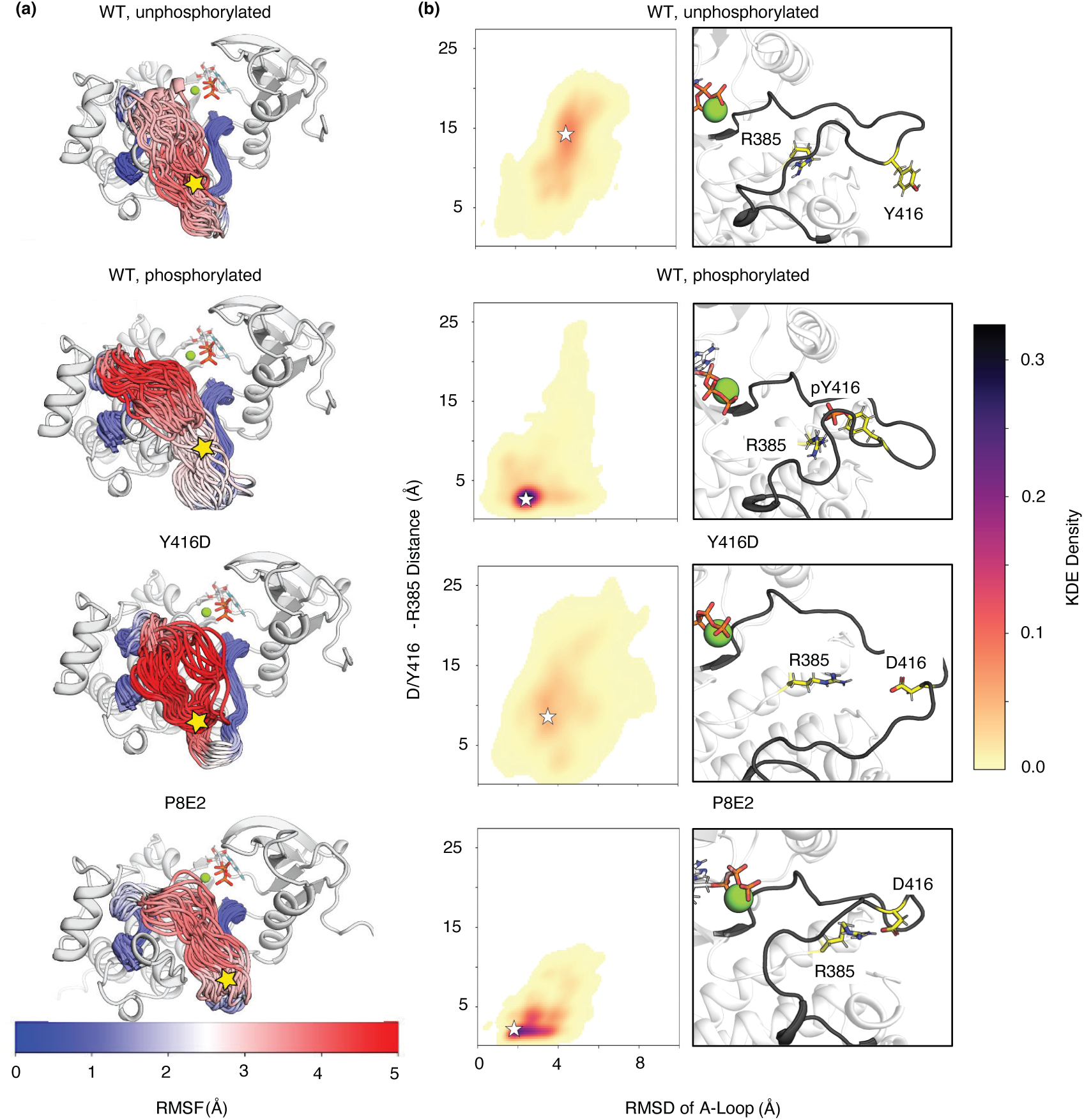
MD Simulations Reveal How Activation Loop Flexibility Differs Between WT, P8E2, and Y416D SRC Kinase Variants. **(a)** Root mean square fluctuation (RMSF, Å) analysis of the activation loop (A-loop) C_α_-atoms was projected onto representative frames from the simulations, where residue Y/D416 is marked with a yellow star. **(b)** 2D kernel density estimates (KDE) of the distance between the Y/D416 and R385 side chains versus the root mean square deviation (RMSD, Å) of the A-loop. The shortest distance was plotted between the terminal nitrogen atoms of the R385 side chain to the oxygen atoms of the D416 or phosphate group. The RMSD was calculated for the backbone atoms of the A-loop with the reference to the starting structure of each system. The representative frames of the highest density region were extracted to illustrate the dominant conformation and are marked on the plots with white stars.

To probe the effects of A-loop deletions on the activation dynamics, we generated 50 diverse AF2 conformers per system that were used to seed ensemble MD simulations (Figure S1). Each simulation was carried out for 400 ns to capture local A-loop and αC-helix rearrangements. The conformational changes were compared in terms of catalytically relevant descriptors: a difference in the distances between residues E310-R409 and K295-E310, and in the angle between the Cα-atoms of K295-E310-Y/D416. Shortening of the distance between K295 and E310 implies a αC-helix rearrangement and formation of a catalytic salt bridge, indicative of an active state. In contrast, a decrease in the E310-R409 distance is associated with the formation of the electrostatic contact that secures the A-loop in the retracted, inactive state (Figure S1). Additionally, the angle between K295-E310-Y/D416 provides an additional description of loop extension and tilt.

Simulation of WT SRC kinase illustrates the formation of a conventional inactive state, with an ordered retracted conformation of the A-loop and its progression towards the extended loop conformation and active retracted αC-helix via a narrow range of intermediates (Figure 3a). In contrast, deletions of P8E2 disrupt the formation of two ordered helical motifs within the loop and destabilize formation of the conventional inactive state. Instead, the inactive state is replaced by a range of retracted loop minima facilitated by the electronic environment of R409 (such as the R409-D413 interaction or proximity of ATP phosphate group), providing a variety of pathways for further formation of the active state (Figure 3b). Additionally, based on the analogous simulations of WT-T338I, P8E2-T338I and P8E2-T338M mutants, the population of these intermediates is readily shifted towards an active state (Extended Figure 3). In the case of P8E2-T338M, the stabilized distance between the E310 and R409 is even greater than in the case of WT or the WT-T338I mutant (displayed by increased population density at the Distance Difference > 6.5 Å), which correlated with an improved activity of this mutant (Extended Figure 1c and Table 1).

**Figure 3:**
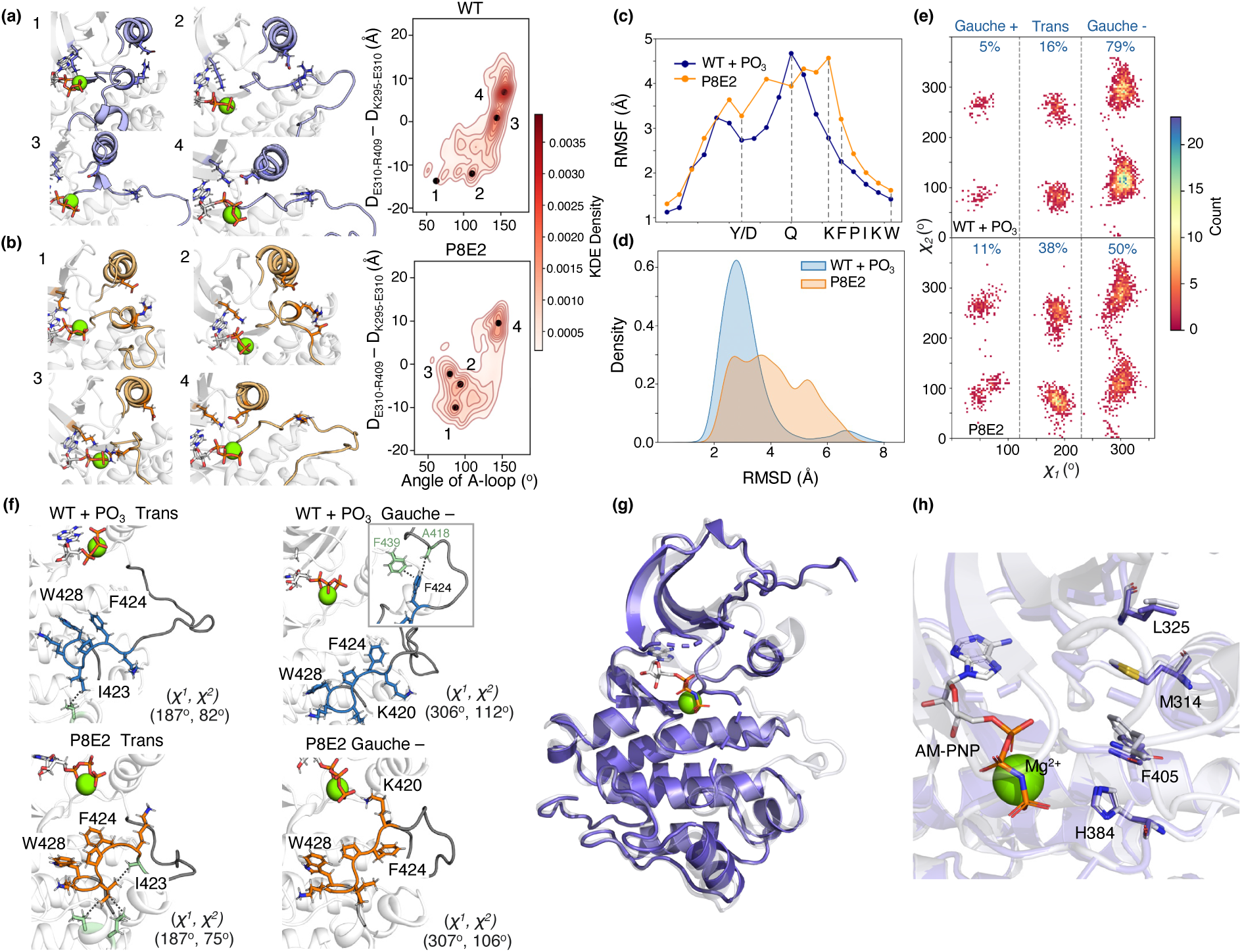
Activation Loop Deletions Reshape SRC Kinase Conformations and Structural Dynamics. Kernel density estimate (KDE) plot and visualization of stabilized minima from combined data from simulations initialized for AlphaFold2^63^ (AF)-seeded MD simulations of (**a**) WT SRC kinase and (**b**) the P8E2 variant. Conformations were analyzed in terms of the salt bridge distances between the side chains of E310-R409 and K295-E310, and the angle between the C_α_-atoms of K295-E310-Y416 for WT SRC kinase, and K295-E310-D416 for the P8E2 variant. (**c**) C_α_-atom root mean square fluctuations (RMSF, Å) for the WT pY416 (blue) and P8E2 (orange) variants, calculated with reference to the initial structure used for each set of simulations. A noticeable shift in the residue mobility towards the KFPIKW segment is observed in the P8E2 variant, with K423 showing the highest RMSF. (**d**) Backbone RMSD distributions of the KFPIKW shelf referenced to the initial structure. The WT A-loop oscillates narrowly around 2– 3 Å from the reference state, while the shortened loop in P8E2 accesses a broader, higher RMSD ensemble relative to the reference state. **(e**) Two-dimensional χ_1_/χ_2_ histogram plots for the F424 side chain (number of bins used = 100). The WT ensemble is dominated by a gauche⁻ rotamer (χ_1_≈ 300°, 79 % of simulation frames) that exposes the phenyl face for π-stacking, while trans rotamers that bury the ring are rare (16 %). In the P8E2 variant, the gauche⁻ population drops to 50 % and the trans state rises to 38% of simulation frames. Percentages were calculated based on the percent of simulation frames that lie in each respective conformation. (**f**) Representative snapshots from the highest density regions of trans (left) and a gauche⁻ (right) F424 rotamers in WT (top) and P8E2 (bottom) SRC kinase. **(g)** X-ray crystal structure of P8E2 (dark purple, PDB ID: 9V4E, this work) at 1.78 Å resolution overlaid with the structure of WT-T338I (lavender, PDB ID: 3DQW^37^). **(h)** Overlay of the hydrophobic spine highlights an intact spine in P8E2, consistent with the active conformation.

MD simulations and structural analysis revealed that the P8E2 deletions also reshape the P+1 loop, particularly the K423–W428 segment, repositioning it toward the catalytic cleft. Increased loop dynamics of this region result in a change of the cleft size and positioning of the aromatic residues linked to substrate selectivity (Figure 3c-e). In WT pY416, residues F424 and W428 form a planar aromatic platform that favors peptide substrates with aromatic residues at the P+3 position (Figure 3f). In P8E2, the shortened loop caps over F424, burying its π-face and creating a narrow hydrophobic groove, consistent with the experimentally observed shift toward aliphatic P+3 substrates (Figure 3e and 3f). These findings demonstrate that loop remodeling can fine-tune substrate recognition by altering the local conformational landscape and side-chain orientations critical for peptide binding.

The dynamic features of the P8E2 variant predicted by simulations are reflected in crystallographic data: the A-loop is largely unresolved, and regions above and below exhibit elevated B-factors (Figure S2), while retaining an intact hydrophobic spine characteristic of the active kinase conformation (Figures 3g and 3h). Together, these results indicate that the triple deletion increases the conformational space sampled by the loop, replacing a well-ordered inactive state with a continuum of dynamic intermediates. This structural flexibility accelerates A-loop opening, improving catalytic efficiency and subtly reprogramming substrate specificity without disrupting the overall kinase fold.

### Activation Loop Deletion in SRC Kinase Modulates Cellular Phosphorylation Profiles and Substrate Preferences

To evaluate how activation loop modifications impact kinase function in a cellular context, we performed unbiased phosphoproteome profiling of HEK293T cells expressing WT SRC, P8E2, and gatekeeper mutants. Principal component analysis (PCA) revealed clear separation of experimental groups along PC1, indicating variant-specific global phosphorylation patterns (Figure S3). Across all conditions, 1,682 differentially expressed (DE) proteins and 5,767 DE peptides were identified (adjusted p < 0.05, |log_2_ fold change| > 1.0), out of 6,419 proteins and 33,127 peptides initially detected (Table S2). Notably, SRC itself was found to be among the most significantly DE proteins when comparing the GFP control versus WT and P8E2, with phosphorylation changes extending beyond the deleted A-loop Y416 site in P8E2, likely reflecting autophosphorylation at alternative sites or activation of endogenous SRC family members (Figures 4a and 4b; Extended Figures 4a and 4b).

**Figure 4:**
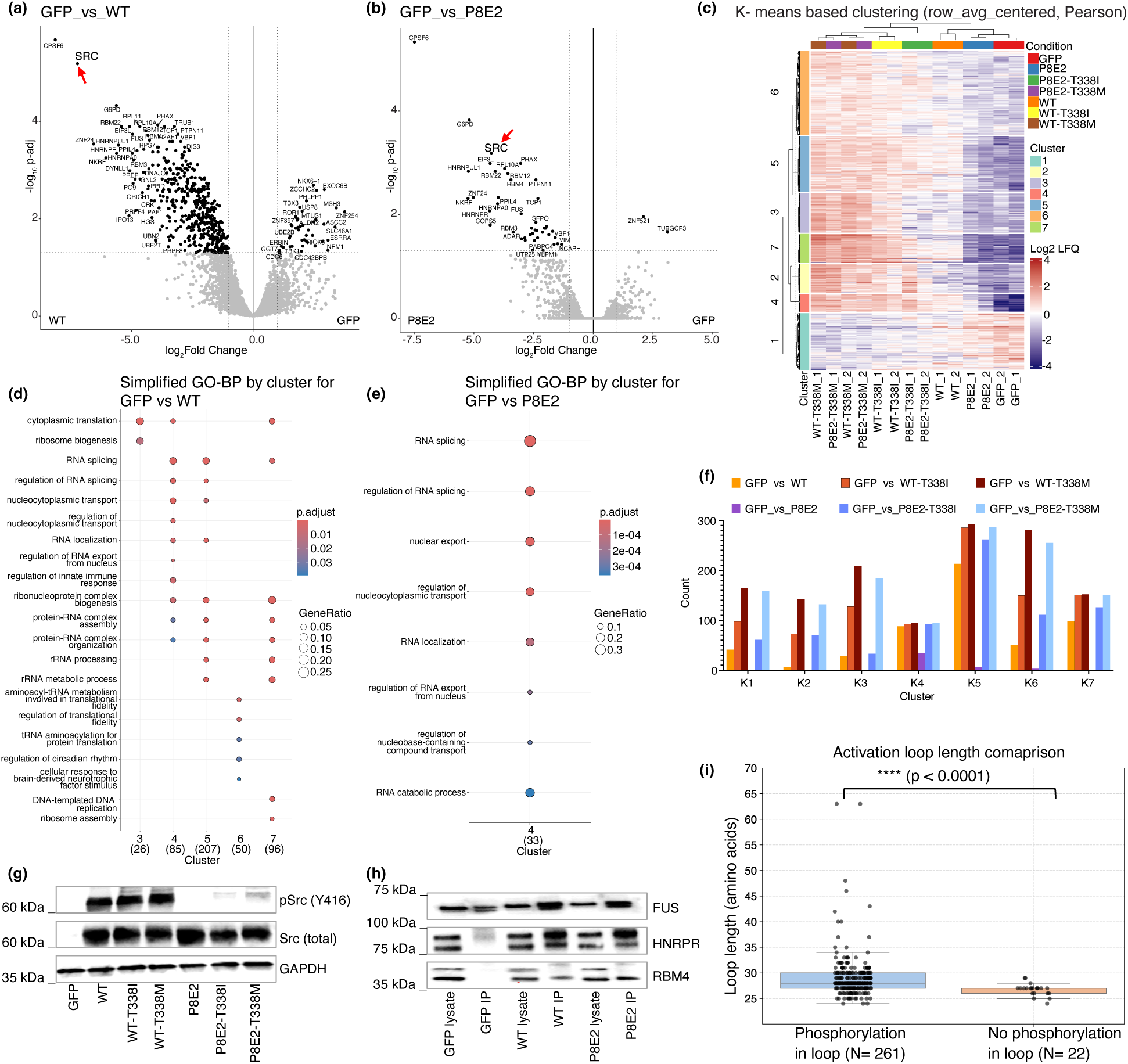
Activation Loop Deletions Rewire SRC Kinase Signaling Networks Revealed by Quantitative Proteomics. **(a and b)** Volcano plots show that the WT SRC kinase domain **(a)** and the P8E2 variant **(b)** induce differential expression of proteins (closed dots) compared to the GFP control. Gray dots represent proteins not considered significant. Vertical and horizontal dotted lines indicate the cutoff line of the magnitude of log2 fold change (−1.0 or 1.0) and FDR (Benjamini-Hochberg) of 0.05, respectively. **(c)** Hierarchical heatmap shows that DE proteins are grouped well by K-means-based clusters for their distributions across experimental conditions. The number of K-clusters was arbitrarily set to seven. The heatmap was row-averages-centered and assessed with Pearson correlation. **(d and e)** Enrichment analysis of DE proteins grouped by the K-means clusters for Gene Ontology Biological Process. WT SRC kinase domain-induced DE proteins are significantly enriched in K-means clusters K3-K7 **(d)**, while P8E2 induced significant enrichment only in cluster K4 **(e)**. **(f)** Bar graph indicates the number of DE proteins in each comparison grouped by K-means clusters (K1-K7). The gatekeeper mutation T338M induced a greater number of DE proteins than that of T338I in K1-K3 and K6 but not in K4, K5, nor K7 when included in the WT or P8E2 SRC kinase domain (FDR < 1 × 10^−4^). Both gatekeeper mutants increased the number of DE proteins greater than the base WT or P8E2 except that in K4, they promoted only P8E2 significantly but not WT (FDR < 1 × 10^−6^). While WT and the gatekeeper mutations have similar effects on the number of DE proteins in K4, it is noteworthy that this cluster was the most significant K-means cluster for P8E2 among all clusters (FDR < 1 × 10^−4^). WT SRC induced DE proteins greatest in K5, followed by K7 and K4 (FDR < 1 × 10^−4^). **(g)** Immunoblot shows that autophosphorylation of SRC Y416 (Y419 in human SRC) is readily detectable in cells transfected with SRC WT, WT-T338I or WT-T338M variants. The P8E2 mutation lacks this tyrosine residue, yet P8E2 mutants compounded with T338I or T338M induce detectable levels of proteins reactive to the phosphoY416 antibody. Anti-SRC (C-terminal) antibody detects overexpression of SRC in the transfected cells except for the GFP control, while anti-GAPDH shows a loading control. WT SRC kinase domain and its variants are fused with GFP. **(h)** Both WT and P8E2 induce tyrosine phosphorylation in DE proteins identified in K-means cluster K4 in transfected cells. Three such proteins, FUS, HNRPR, and RMB4, are shown. Endogenous proteins were enriched in cell lysates by anti-GFP antibody-conjugated beads as overexpressed SRC and its variants are tagged/fused with GFP proteins. Tyrosine-phosphorylated proteins were immunoprecipitated in the enriched lysates using agarose beads conjugated with anti-phosphotyrosine antibody. Immunoprecipitates (IP) were used for immunoblots to probe the K-means cluster K4 proteins in the corresponding lysates. **(i)** Comparison of activation loop lengths (spanning from the DFG to APE motifs) among human kinases. Kinase activation mechanisms and loop phosphorylation status were verified using UniProt, and kinases with unclear mechanisms were excluded from the analysis. Within the “No Phosphorylation in Loop” group, kinases from the CK family displayed a uniform loop length of 35 amino acids and were excluded as outliers. Differences in loop length between the two groups were assessed using a two-tailed Mann–Whitney U test.

Closer examination of A-loop peptides confirmed the absence of Y416 phosphorylation in P8E2, whereas peptides corresponding to conserved sequences in other SRC family members (e.g., LYN, HCK) were differentially phosphorylated in gatekeeper mutant backgrounds, indicating cross-activation of endogenous kinases (Table S3).

K-means clustering of DE proteins and peptides into seven clusters revealed variant-specific patterns of substrate engagement (Figures 4c and 4f, Extended Figures 4c and 4d). WT SRC induced broad changes across multiple clusters, particularly K4, K5, and K7, associated with enrichment terms linked to RNA processing, protein synthesis, and DNA replication. In contrast, P8E2 predominantly influenced a narrower set of targets within K4, enriched for RNA processing and regulation (Figures 4d and 4e). These findings suggest that deletion of N414, T417, and R419 constrains SRC activity to a selective subset of substrates within the shared substrate pool, disproportionally skewed toward cluster K4 (Figure 4f and S4). This restricted engagement parallels the shift in substrate selectivity observed in vitro, together supporting the idea that minimal loop perturbations reprogram kinase specificity.

When the number of DE proteins are plotted for all comparison sets and grouped by K-clusters, clusters K5 and K6 contained a high number of DE proteins in both gatekeeper variants of WT and P8E2 (Figure 4f, Extended Figure 4d), reflecting broad substrate engagement. In contrast, cluster K1 displayed an opposite heatmap pattern for the highly active gatekeeper-containing variants (Figure 4c). Interestingly, DE proteins in this cluster are associated with positive regulation of cell cycle and proliferation (Extended Figures 5a and 5b). EGFR and other positive regulators of proliferation were identified among these GO terms in GFP versus WT-T338M comparison (Extended Figure 5c); therefore, it is tempting to speculate that the hyperactive SRC variant may trigger negative feedback mechanisms to modulate proliferation signaling (Extended Figures 6 and 7). These observations underscore that the P8E2 deletion variant alone enables selective phosphorylation, while addition of gatekeeper mutations drives widespread engagement and compensatory feedback, highlighting the importance of balanced catalytic activity and substrate specificity for controlled signaling.

Western blot validation confirmed high phospho-SRC levels in WT and gatekeeper mutants, while P8E2 variants displayed lower phosphorylation, supporting the proteomics results (Figure 4g). Selected DE targets from the K4 cluster were further validated, revealing that P8E2 induces highly selective phosphorylation within these shared targets at levels equivalent to that of WT (Figure 4h). Together, these data indicate that minimal deletions within the activation loop can reprogram SRC signaling in cells by narrowing substrate engagement and enhancing selective phosphorylation.

### Nature Employs Activation Loop Shortening to Control Kinase Regulation

To investigate whether the length of the activation loop was shaped by evolutionary pressures as a determinant of kinase regulation, we constructed a phylogenetic tree using approximately 400 kinase domain sequences (Supplementary Data 3) from diverse taxa, including Amoebozoa, Fungi, Choanoflagellates, Filasterea, and Metazoa. These lineages span both unicellular and multicellular organisms. As expected, core catalytic motifs such as HRD in the catalytic loop and DFG and APE in the activation loop were highly conserved across all clades (Figure S5), reinforcing the deep structural and functional conservation among tyrosine kinases^41^.

The phylogenetic analysis revealed a clear separation between early-diverging groups (Amoebozoa and Fungi) and later-branching lineages (Choanoflagellates, Filasterea, and Metazoa), with the latter forming a well-supported clade suggestive of a shared evolutionary trajectory (Figure S6). Receptor and non-receptor tyrosine kinases also clustered into distinct clades, reflecting their functional divergence. Notably, canonical autophosphorylation sites were largely absent in Amoebozoa and Fungi, indicating a simpler signaling framework in these primitive eukaryotes (Figure S5, red and blue arrow). In contrast, these regulatory sites became increasingly conserved among Metazoan kinases, appearing in nearly all tyrosine kinases analyzed.

A particularly distinct clade was formed by the C-terminal Src kinase (CSK), a non-receptor tyrosine kinase that lacks an autophosphorylation site and phosphorylates the C-terminal tail of SRC family kinases (Figure S6) with exceptional specificity^42^. Unlike most tyrosine kinases, CSK utilizes acidic residues within its activation loop to mimic the effect of phosphorylation, bypassing the need for canonical activation. Remarkably, CSKs from all species examined possess a significantly shorter activation loop (25 amino acids), a feature that is highly conserved across evolutionary time. Given that both CSK and SRC are among the few tyrosine kinases that evolved prior to the divergence of unicellular and multicellular organisms^43,44^, the conserved short loop length in CSK strongly suggests that activation loop length was already under selective pressure in early eukaryotes. This observation supports the hypothesis that activation loop length served as a predictive determinant of kinase regulatory mode well before the rise of complex multicellular signaling. Thus, CSK represents a compelling evolutionary example of how structural constraints on the activation loop may have been established in pre-metazoan lineages to enable alternative regulatory strategies beyond autophosphorylation.

To investigate whether the effect of activation loop shortening is a broader regulatory strategy among kinases lacking loop phosphorylation, or a unique feature restricted to CSK, we analyzed the activation loop length across all human kinases. We categorized the kinases into two groups: those that require phosphorylation within the activation loop for activation, and those that do not (Supplementary Data 4). Our analysis revealed that approximately 11% of human kinases function without regulatory phosphorylation in the loop. Notably, around 76% of these kinases exhibit a significantly shorter activation loop (Figure 4i), suggesting that loop shortening may represent a conserved alternative mechanism for achieving kinase regulation in the absence of canonical activation loop phosphorylation. This dual mechanism, combining the absence of an autophosphorylation site with a structurally constrained loop may represent a broader evolutionary strategy by which certain kinases attain precise substrate recognition. Our engineered deletion variants may have recapitulated aspects of this strategy, suggesting that similar mechanisms can be engineered to rewire kinase specificity.

## Discussion

Protein kinases form finely tuned signaling networks in which the coordinated action of multiple enzymes and mediators discriminates among substrates to orchestrate complex cellular responses^45,46^. While structural and sequence analyses have identified determinants of substrate specificity^47^, translating this knowledge into strategies to reprogram kinase signaling has remained underexplored. Traditional approaches, such as ATP-pocket engineering, primarily facilitate target mapping rather than modulating intrinsic specificity^48,49^. Here, we focused on the activation loop, a dynamic and evolutionarily variable element, as a lever to redirect kinase function. Our results show that minimal, localized modifications within this loop can reshape kinase behavior: substitution of the autophosphorylation site Y416 with aspartate impaired catalytic efficiency, whereas a targeted triple-deletion variant (P8E2) restored activity and subtly shifted substrate preference toward acidic residues. These effects were amplified in cellular contexts, demonstrating how small structural adjustments can propagate through kinase networks to rewire signaling outcomes.

Insights from nature underscore this principle. For instance, structural analyses of CSK revealed that deletion of specific anchor residues in its activation loop underlies elevated specificity toward SRC-family kinases. Yet, deletion of analogous residues in SRC caused a tenfold reduction in catalytic activity, suggesting that loop length and composition encode more sophisticated rules of substrate selection than simple truncation alone^42^. Our work provides a direct test of this idea, showing that rational, minimal changes within the activation loop can recover activity while redirecting specificity.

This strategy redefines the current paradigm of synthetic network design, which has largely relied on orthogonal interaction domains^16^. While such approaches offer predictability and modularity, they remain disconnected from the evolutionary principles and molecular logic that govern natural kinase signaling. By contrast, our results reveal that the activation loop encodes a compact regulatory code that can be rationally reprogrammed. Rather than constructing entirely new circuits, this approach enables kinases to function within their native signaling environment while selectively rewiring specific connections. This provides a path to rewire existing nodes of cellular networks and uncovers how regulatory loop features shape distinct biological outputs.

Our findings also address a long-standing uncertainty in the field: whether substrate preference is distributed across the entire kinase domain or localized to modular regulatory elements. By showing that targeted loop changes can redirect signaling outcomes without abolishing catalytic activity, we demonstrate that the activation loop itself encodes substrate discrimination logic. The broader implications extend beyond fundamental biochemistry. In synthetic biology, this approach offers a scalable strategy to create kinases that reprogram signaling cascades by redirecting endogenous nodes rather than adding orthogonal ones, opening a new paradigm for both understanding and engineering cellular networks.

To test whether this principle extends beyond SRC, we introduced equivalent mutations into two additional SRC-family kinases, Hck and Fyn. In these cases, mutation of the conserved autophosphorylation site was more deleterious than in SRC, yet analogous deletions successfully restored catalytic activity (data not shown). These results suggest that a kinase’s ability to function without its canonical regulatory switch is strongly correlated with activation loop length, a feature not previously recognized. Ongoing work is aimed at mapping the distinct signaling responses that arise from these variants in cellular contexts.

Together, these findings highlight loops as uniquely powerful elements for precision engineering. Unlike rigid α-helices or β-strands, loops exhibit high sequence and structural variability, act as dynamic connectors, and strongly influence enzyme flexibility^50–52^. Exploiting insertions and deletions (InDels) within these regions produces functional shifts^53^ that substitutions alone cannot achieve, opening a new dimension in protein engineering. While most experiments here were performed with isolated kinase domains, our results establish a conceptual framework for manipulating full-length kinases and rewiring complex signaling networks. Future studies in cellular and organismal models will be essential to validate and expand these insights.

In conclusion, we demonstrate that minimal, localized modifications within the activation loop can systematically reprogram kinase activity and substrate specificity. This establishes a generalizable strategy for designing programmable kinase circuits, providing precise control over signaling outcomes and advancing both fundamental kinase biology and synthetic applications.

## Materials and Methods

### Cloning and Mutagenesis of Chick SRC Kinase Domain and YopH Phosphatase

The codon-optimized genes encoding the chick SRC kinase domain (UniProt ID: P00523, residues 251–533) and YopH phosphatase (UniProt ID: P15273) were synthesized by TWIST Bioscience for expression in *E. coli*. The SRC gene was cloned into the pET-28a(+) vector with an N-terminal His tag, while the YopH gene was cloned into the pCDFDuet-1 vector, using In-Fusion Cloning (Takara Bio). Additionally, the genes for WT SRC and the P8E2 mutant (amplified from the library) were cloned into the pCDNA3.1 vector, with an N-terminal eGFP tag, for expression in HEK293T cells following the same In-Fusion Cloning protocol.

PCR amplification of the synthetic genes and linearized vectors was carried out using PrimeSTAR Max DNA polymerase (Takara Bio) with primers containing appropriate overlapping regions. Site-directed mutagenesis was performed via inverse PCR using the same polymerase and specifically designed primers (Supplementary Data 2). The accuracy of cloning and mutagenesis was verified by Sanger sequencing.

### Multiple Sequence Alignment for SRC and Related Kinases

To identify residues that differ from the consensus sequence in the activation loop of SRC, a BLAST search was performed using the human SRC sequence (UniProt ID: P12931), retrieving 5,000 hits. Redundant sequences were removed using CD-HIT^54^ with a 90% sequence identity cutoff, resulting in 496 unique sequences (Supplementary Data 1). These sequences were aligned using the MUSCLE^55^ alignment tool. No additional filtering was applied. Residues corresponding to positions 407–435 (activation loop, based on human SRC numbering) were manually inspected, and the identified divergent positions (Table S1, Supplementary Data 2) were used to design the mutant library.

### Substitution Library

A codon-optimized gene library was designed using a 150 bp megaprimer to introduce substitutions in the activation loop of SRC. The megaprimer was selected within the region spanning amino acids 391–442 and encoded all theoretically possible 131 single and double mutants (Supplementary Data 2). The library was synthesized as an oPools™ Oligo Pool by Integrated DNA Technologies (IDT).

PCR amplification of the library was performed using PrimeSTAR Max DNA polymerase. The amplified library was cloned into the pET-28a(+) vector using a megaprimer approach and whole-plasmid cloning (MEGAWHOP)^56^. A touchdown PCR^57^ protocol was employed, gradually decreasing the annealing temperature from 72°C to 47°C by 1°C per cycle. The resulting cloning product was transformed into *E. coli* DH5α chemically competent cells (SMOBIO Technology Inc).

Library diversity was assessed by randomly picking and sequencing several colonies, confirming broad coverage. To isolate the plasmids from the transformed library, all colonies from the agar plate (several thousand, kanamycin-resistant) were scraped and pooled. Plasmids were then extracted from the pooled *E. coli* pellet using the NucleoSpin^®^ Plasmid EasyPure mini prep kit (Macherey-Nagel).

### Deletion Library

Following a similar approach to the substitution library, a codon-optimized gene library was designed for deletions spanning amino acids 389–445. This library was obtained as an oPools™ Oligo Pool from IDT and cloned using In-Fusion Cloning (Takara Bio).

PCR amplification of the library and vector was performed using PrimeSTAR Max DNA polymerase with primers containing slightly longer overlaps. The cloning product was then transformed into *E. coli* DH5α chemically competent cells. Plasmid isolation for the deletion library followed the same protocol as the substitution library.

### Large Scale Protein Expression and Purification

Protein expression and purification for SRC and its variants were carried out using a previously adapted protocol^58^. Briefly, a single colony of *E. coli* BL21 (DE3) (SMOBIO Technology Inc). co-transformed with kinase (pET-28a (+)) and phosphatase (pCDFDuet-1) plasmids was picked from an agar plate and inoculated into LB medium supplemented with kanamycin (50 μg/ml) and streptomycin (50 μg/ml). The culture was incubated overnight at 37°C with shaking at 220 rpm. The following day, the overnight culture was diluted into TB medium containing the same antibiotics and grown at 37°C with shaking at 220 rpm until the OD_600_ reached 0.8–1.0. The culture was then cooled, and protein expression induced by adding IPTG to a final concentration of 0.2 mM, followed by incubation at 18 °C with shaking at 220 rpm for at least 16 h. Cells were harvested by centrifugation at 4,000 rpm for 20 min at 10 °C, and the resulting pellet either stored at −80 °C or used directly for protein purification.

#### Purification

Protein purification was carried out using a two-step process: His-tag purification via Ni-NTA affinity chromatography followed by size exclusion chromatography (SEC). For His-tag purification, *E. coli* cell pellets were thawed and resuspended in lysis buffer (50 mM Tris pH 8.0, 500 mM NaCl, 5% (v/v) glycerol, 25 mM imidazole) supplemented with benzonase (25 U/ml) and lysozyme added directly as a solid. Cells were lysed using sonication or microfluidization, and the lysate was clarified by centrifugation at 10,000 rpm for 30–40 min at 10 °C. The supernatant was applied to His60 Ni Superflow resin (#Z5660N, Takara Bio), pre-equilibrated with lysis buffer, and washed extensively to remove non-specifically bound proteins. The bound protein was eluted in fractions using the same buffer containing 0.5 M imidazole (Buffer B). Elution fractions were analyzed by SDS-PAGE (4-15 % Mini-PROTEAN TGX, BioRad) to confirm protein purity. Following elution, the protein was directly loaded onto a pre-equilibrated size exclusion chromatography column and eluted using exchange buffer (50 mM Tris pH 8.0, 100 mM NaCl, 5% (v/v) glycerol, 1 mM TCEP). Protein concentration was evaluated spectrophotometrically using the molar absorption coefficient calculated in ProtParam.

### Replacing the Thrombin Site with a TEV Site in the P8E2 Vector

The Thrombin site in the pET-28a(+) vector containing P8E2 was replaced with a TEV site in a single step using inverse PCR. The reaction was performed with PrimeSTAR Max DNA polymerase and specifically designed primers.

### TEV Protease Expression and Purification

The plasmid encoding MBP-fused TEV protease, which includes a self-cleaving TEV site positioned between MBP and the 7×His tag, was generously provided by the Colin Jackson lab. During expression, the TEV site undergoes cleavage, separating MBP from the His-tagged TEV protease. TEV protease was expressed and purified following the same protocol and buffer conditions used for SRC variant expression (Figure S7 and S8).

### His-Tag Removal and Purification

Purified His-tagged kinase P8E2 with TEV cleavage site was mixed with TEV protease at a 1:10 molar ratio of TEV protease to kinase in exchange buffer and incubated overnight at 10 °C without shaking. Cleavage efficiency was monitored the following day by SDS-PAGE, which showed over 99% cleavage. To remove the TEV protease from the mixture, the solution was applied to His60 Ni Superflow resin pre-equilibrated with exchange buffer. The flow-through was passed through the resin 5-6 times to maximize binding of the TEV protease. To remove any nonspecifically bound kinase, the resin was washed with exchange buffer, and fractions of 1 mL were collected. Fractions containing cleaved protein were pooled and further purified using SEC with the same exchange buffer (Figure S7 and S8).

### X-ray Crystallography

Crystals of P8E2 without His-tag were grown at 10 °C using the hanging drop method. P8E2 was prepared at a concentration of 6-7 mg/mL in exchange buffer and mixed with AMPPNP and MgCl_2_ to achieve final concentrations of 5 mM and 25 mM, respectively, resulting in a homogeneous solution. 2 μL drops were set up, each containing 1 μL of protein + cofactors and 1 μL of reservoir solution, which were equilibrated over 500 μL of reservoir solution. The reservoir conditions were screened between 15-20% PEG 3350 with 5% (v/v) glycerol, 0.1 M Bis-Tris (pH 5.5-7.0), and 0.2 M sodium acetate. Crystals formed during this screening were used to generate seeds with the Seed Bead Steel Kit (HR4-780) from Hampton Research. The microseeds, diluted serially, were then used in a new crystallization screen with the same reservoir conditions, varying only the PEG 3350 concentration between 15-20% (w/v) at pH 5.5. In the new drops, 1 μL of protein + cofactor was mixed with 1 μL of microseed at different dilutions. Finally, crystals were obtained under the conditions of 19% (w/v) PEG 3350, 5% (v/v) glycerol, 0.1 M Bis-Tris pH 5.5, and 0.2 M sodium acetate. All optimization steps and final crystallization were performed at 10 °C. The crystals were cryoprotected with 30% (v/v) PEG 400 in mother liquor, flash-frozen, and stored in liquid nitrogen.

X-ray diffraction data were collected at the SPring-8 synchrotron (Harima, Japan) on beamline BL45XU with an X-ray wavelength of 1.00 Å and a temperature of 100 K. Automated data collection was performed using the ZOO suite^59^, and the data were automatically indexed, integrated, scaled and merged using XDS^60^ accessed via the KAMO pipeline^61^. The structure was solved by molecular replacement in MOLREP^62^ using an AlphaFold2 model as a search model^63^. The structure was refined by iterative real-space and reciprocal-space refinement in COOT^64^ and REFMAC5^65^, respectively. Data collection and refinement statistics are given in Supplementary Table 4.

### Library Expression

The isolated library from *E. coli* DH5α cells was co-transformed with YopH into electrocompetent *E. coli* BL21 (DE3). To increase the number of colonies, antibiotic concentrations were reduced to 20 µg/mL kanamycin and 10 µg/mL streptomycin. Individual colonies were picked and inoculated into 96-deep-well plates containing 1 mL LB medium supplemented with antibiotics. The plates were incubated overnight at 37 °C with shaking at 2,000 rpm on a plate shaker.

The following day, ~10 µL of the overnight culture from each well was transferred into a fresh 96-deep-well plate (maximum volume 2 mL) containing ~400 µL TB medium with the same antibiotics and grown at 37 °C with shaking for 3.5 to 4 h. The plates were then cooled, and protein expression was induced by adding IPTG to a final concentration of 0.2 mM, followed by incubation at 18 °C with shaking at 2,000 rpm for at least 16 h. Cells were harvested by centrifugation at 3,700 rpm for 20 min at 10 °C, and the resulting pellets in deep well plates were either stored at −80 °C or used directly for purification. For comprehensive library coverage, ~500 colonies from the substitution library (P1 to P5) and ~300 colonies from the deletion library (P6 to P8) were transformed and screened separately.

### Purification of library variants

Cell pellets were resuspended in 200 µL of 1X BugBuster (#70921, Merck) solution diluted in lysis buffer, supplemented with lysozyme and benzonase, and incubated with shaking at ~1,000 rpm for 20–30 min. The lysate was clarified by centrifugation at 3,700 rpm for 20 min at 10 °C. Meanwhile, 96-deep-well filtration plates (1 mL, 50 μm Frit) charged with His60 Ni Superflow resin, were prepared. Approximately 200 µL of resin was aliquoted into each well and equilibrated with lysis buffer. The filter plates were attached to standard 96-deep-well collection plates (maximum volume 2 mL) and centrifuged at 500 × g for 2 min.

The clarified lysate was transferred to the filter plates and centrifuged once to collect the flowthrough. The flowthrough was then reapplied to the resin to maximize protein recovery. The resin was washed four times with ~500 µL of lysis buffer to remove nonspecifically bound proteins. Finally, the bound protein was eluted in 100 µL of Buffer B. Eluted fractions were used directly for kinase activity screening.

### Enzymatic Assay

A classical coupled enzymatic assay utilizing pyruvate kinase (#P9136, Sigma-Aldrich) and lactate dehydrogenase (LDH, #L1254, Sigma-Aldrich) was used to monitor the phosphorylation activity of SRC variants^58^. The assay was performed in a UV-transparent 96-well plate (# 2145-40, funakoshi), where the rate of phosphorylation of the peptide substrate (EAIYAAPFAKKK; GL Biochem Ltd.) by the kinase of interest, measured through ATP (# A7699, Sigma-Aldrich) consumption, was coupled to the oxidation of NADH (# 10128023001, Sigma-Aldrich) by LDH. The reaction was carried out in 100 mM Tris buffer (pH 8.0) with final concentrations of 0.4 µM LDH, 0.2 µM pyruvate kinase, 1 mM phosphoenolpyruvate (#P7127, Sigma-Aldrich), 0.2 mM peptide substrate, 10 mM MgCl_2_, and 0.2 mg/mL NADH. NADH consumption was continuously monitored at 340 nm for 30 min.

### Library Screening

For high-throughput screening of the SRC variant library, the coupled enzymatic assay was performed at a fixed ATP concentration of 2 mM. A reaction cocktail containing all reagents for a 200 µL final volume was prepared, and the volume was adjusted to 160 µL with water before enzyme addition. Then, 40 µL of eluted protein fractions from the 96-well expression plates was directly added to initiate the reaction.

The reaction rate was determined by monitoring NADH consumption at 340 nm for 30 min, with readings taken every 6 s, and the rate of decrease within the linear range was calculated using Prism software. Variants displaying at least 65% of the WT reaction rate were selected and sequenced. A total of eight plates were analyzed—five from the substitution library and three from the deletion library. From these, 60 variants were selected, pooled, expressed, purified, and reanalyzed together in a single plate. Based on this reanalysis, the top five mutants were selected for large-scale purification and kinetic characterization.

### Kinetic Analysis

Michaelis–Menten kinetic parameters for selected SRC variants were determined using the same coupled enzymatic assay. ATP concentrations were varied as follows: 0, 0.04, 0.08, 0.16, 0.32, 0.65, 1.25, and 2.5 mM. An additional ATP concentration of 0.02 mM was included for gatekeeper variants. Reactions were initiated by adding either 0.2 µM kinase or 0.1 µM for gatekeeper variants. The final reaction volume, including the kinase, was 200 µL, with water used to adjust the volume before enzyme addition. Continuous NADH consumption was monitored at 340 nm for 30 min, with readings taken every 6 s. Kinetic parameters were determined by fitting the data to the Michaelis–Menten equation using Prism software.

### Peptide Library Expression for High-Throughput Substrate Specificity Screening Assay

To determine substrate specificity, a high-throughput screening assay developed by Kuriyan et al^38,39^ was followed with slight modifications to enzyme concentration and reaction time.

The Human-pTyr library (from the Kuriyan lab) was transformed into *E. coli* MC1061 electrocompetent cells. Following recovery, 100 µL of the transformed cells were incubated in 5 mL of fresh LB medium supplemented with chloramphenicol (25 μg/ml) and grown overnight at 37°C with shaking at 220 rpm. The next day, 100 µL of the overnight culture was diluted into fresh 5 mL LB medium containing antibiotics and grown until the OD_600_ reached 0.5–0.6. At this point, peptide library expression was induced with 0.4% (w/v) arabinose (final concentration) for 4 h at 25 °C. After induction, cells were harvested in small aliquots (~700 µL) and washed with PBS before proceeding with cell surface phosphorylation.

### Cell Surface Phosphorylation

Phosphorylation reactions were carried out at 37 °C for 3 min, with kinase concentrations optimized to achieve approximately 20–25% cell surface phosphorylation. The kinase concentrations used were as follows: WT and P8E2, 100 nM; WT-T338I, P8E2-T338I, and WT-T338M, 25 nM; P8E2-T338M, 10 nM. A premix of kinase and reaction buffer containing 50 mM Tris pH 7.5, 150 mM NaCl, 5 mM MgCl_2_, 1 mM TCEP, and 2 mM sodium orthovanadate was prepared, and cell pellets were resuspended in this premix. Reactions were initiated by adding ATP to a final concentration of 1 mM and quenched by the addition of 500 mM EDTA pre-incubated on ice.

Cells were harvested by centrifugation at 4,000 rpm, washed twice with binding buffer (PBS + 0.2% (w/v) BSA), and labeled with the pan-phosphotyrosine antibody 4G10-PE (#FCMAB323PE, Merck Millipore) at a 1:100 dilution in binding buffer. After labeling, cells were washed again and subjected to fluorescence-activated cell sorting (FACS) (Figure S9).

### Post-FACS Processing

Sample processing, preparation for deep sequencing, and data analysis were performed as previously described^38,39^ without modifications.

### Cell Culture and Transfection for Proteomics Experiments

HEK293T cells were cultured and maintained at 37 °C in DMEM high-glucose medium (ThermoFischer Scientific) supplemented with 10% fetal bovine serum (#F0926, Sigma-Aldrich). For transfection of eGFP-tagged SRC variant plasmids, HEK293T cells were seeded in six-well plates to approximately 70% confluency. Transfections were carried out using the Lipofectamine 3000 Transfection Reagent following the manufacturer’s protocol. A plasmid encoding only eGFP was used as a control. Transfection efficiency was confirmed 24 hours post-transfection using a fluorescence microscope (Figure S10).

### MS Sample Preparation

Upon transfection cells were trypsinized, collected in 1.5 mL tubes, and prepared for mass spectrometry (MS) analysis. Protein digestion, tryptic peptide purification, and phosphopeptide enrichment were performed using the EasyPep™ Mini MS Sample Prep Kit (#A40006) and the High-Select™ TiO_2_ Phosphopeptide Enrichment Kit (#A32993) from Thermo Scientific according to the manufacturer’s instructions.

### LC-MS/MS Analysis

LC-MS/MS analysis was performed using a nanoElute2 system (Bruker Daltonics) coupled to a timsTOF HT mass spectrometer (Bruker Daltonics) equipped with a CaptiveSpray 2 nano-electrospray ion source (Bruker Daltonics). Peptides were loaded onto an Aurora Ultimate C18 emitter column (25 cm × 75 µm inner diameter, 1.7 µm particle size; ionopticks) fitted with a CaptiveSpray insert. Peptide separation was achieved using a 60-minute linear gradient from 2% to 35% solvent B, at a flow rate of 250 nL/min. Solvent A consisted of 0.1% formic acid in water, and solvent B contained 0.1% formic acid in acetonitrile. The column was maintained at 50 °C using an external column heater (Toaster M, Bruker Daltonics).

Mass spectrometric data were acquired in data-dependent PASEF (Parallel Accumulation–Serial Fragmentation) mode. Each acquisition cycle included one TIMS-MS survey scan followed by ten PASEF MS/MS scans. Ion accumulation and ramp times in the dual TIMS analyzer were set to 100 ms each. The ion mobility range was set from 1/K_0_ = 0.85 to 1.3 V·s/cm². Precursor ions were scanned in the m/z range of 100–1700, with quadrupole switching synchronized to the elution profile from the TIMS device. Collision energy was adjusted linearly based on ion mobility, decreasing from 59 eV at 1/K_0_ = 1.6 V·s/cm² to 20 eV at 1/K_0_ = 0.6 V·s/cm². Singly charged precursors were excluded using a polygon filter. MS/MS precursor selection was based on an intensity threshold of 2,500, with repeated sampling until a target intensity of 20,000 was reached. Dynamic exclusion was applied with a retention time window of 0.4 min.

All raw LC-MS/MS data have been deposited in the ProteomeXchange Consortium via the jPOST repository^66^ under the dataset identifier PXD068232.

### Phosphoproteomic Data Analysis

Raw data were processed in the FragPipe software (v22.0) suite^67,68^. Database searches were performed with MSFragger against the UniProt/Swiss-Prot *Homo sapiens* reference proteome (20,375 sequences, release 2021-04). Search parameters included a precursor mass tolerance of ±20 ppm and fragment ion mass tolerance of ±20 ppm. Enzyme specificity was set to trypsin, allowing up to two missed cleavages. Carbamidomethylation of cysteine residues was set as a fixed modification, while phosphorylation of serine, threonine, and tyrosine was specified as a variable modification. Peptide-spectrum matches were filtered to a 1% false discovery rate (FDR) at the peptide level using Percolator. Label-free quantification (LFQ) was performed with IonQuant using the MaxLFQ algorithm to obtain normalized protein and peptide intensities. The resulting MaxLFQ outputs were then analyzed at both protein and peptide levels using the R package FragPipeAnalystR^69^. As a layer of normalization, ‘vsn’ (variance stabilization normalization) was applied to the proteome data when indicated^70,71^. Missing value imputation was performed with the Bayesian principal component analysis ‘bpca’ option as one of the most versatile and reliable missing value estimation methods for label-free proteome analysis^72,73^. Differential expression (DE) was estimated by the empirical Bayes statistical package ‘limma’ with the default FDR/BH cutoff of 0.05 and fold change greater than 2-fold or smaller than 1/2 in each comparison after conversion of LFQ intensity to log2-scaled values^74^. The protein DE level was further bioinformatically analyzed for enrichment using the R package clusterProfiler version 4.16^75^ with Gene Ontology (Biological Process, Molecular Function, and Cellular Component)^76^. K-means based clustering was used in hierarchical heatmap using the R package ‘ComplexHeatmap’ based on row-average centered and Pearson correlation^77^.

### Statistical Analysis of Differential Protein and Peptide Distributions Across K-Means Clusters

The numbers of DE proteins and peptides within and between K-means clusters were first globally assessed using Monte Carlo simulations applied to a 7 × 6 contingency table (representing seven K-means clusters and six experimental contrast sets). This test evaluated the null hypothesis (H_0_) that DE proteins are randomly distributed across clusters and contrasts. Upon rejection of H_0_ (p = 9.999 × 10^−5^, 10,000 replicates), post-hoc pairwise chi-squared were performed with FDR correction using the Benjamini–Hochberg procedure—either between all combinations of experimental contrasts within each K-means cluster, or between all K-means clusters within each experimental contrast (for example, GFP_vs_WT). All statistical analyses were performed using custom scripts written in the R programming language.

### Protein Extraction for Western Blot Analysis

After transfection, proteins were extracted using ice-cold RIPA buffer supplemented with protease (#87786, ThermoFisher Scientific) and phosphatase inhibitor (#78420, ThermoFisher Scientific) cocktails. Cells were scraped from the culture surface and collected into 1.5 mL microcentrifuge tubes. Lysates were centrifuged at 15,000 rpm for 15 min at 4 °C, and the supernatant was collected. Protein concentration was determined using the Bradford assay.

### Analysis of Src Autophosphorylation

For each sample, 10 µg of total protein was mixed with 4× Laemmli sample buffer containing 100 mM DTT and boiled at 95 °C for 5 min. Proteins were resolved on 4–15 % gradient SDS–PAGE gels at 200 V for approximately 30 min and transferred to PVDF membranes. Membranes were blocked in blocking buffer (Bullet Blocking One; Nacalai Tesque) for 30 min at room temperature, then incubated overnight at 4 °C with primary antibodies against p-Src (Tyr416/419, 1:1000, Cell Signaling Technology, #6943P) and Src(36D10) (1:1000, Cell Signaling Technology, #2109) diluted in blocking buffer. After washing with TBST, membranes were incubated for 1 h at room temperature with HRP-conjugated anti-rabbit IgG secondary antibody (1:5000, Cell Signaling Technology, #7074). Protein bands were visualized using Immobilon Western Chemiluminescent HRP Substrate (#WBKLS0500, Merck) according to the manufacturer’s instructions and detected with an iBright1500 imaging system.

### Analysis of Cluster 4 Proteins by Immunoprecipitation and Western Blot

To detect Cluster 4 proteins (FUS/TLS, HNRPR, and RBM4), lysates containing 500 µg of protein in 1 mg/mL total lysate (adjusted with RIPA buffer) from GFP-alone, WT SRC, and P8E2-expressing cells were first incubated with ChromoTek GFP-Trap® Agarose beads (50 µL) for 1 h at 4 °C to remove overexpressed GFP-tagged proteins. Beads were pelleted by centrifugation at 2,500 × g for 5 min at 4 °C, and the clarified lysates were incubated overnight at 4 °C with 4G10 Platinum Anti-Phosphotyrosine Agarose Conjugate (40 µL). The next day, beads were pelleted at 1,500 × g for 5 min at 4 °C, and the supernatant was discarded. Beads were washed three times with 1× PBS, and bound proteins were eluted by boiling in SDS–PAGE sample buffer at 95 °C for 5 min. Eluted proteins were separated by SDS–PAGE and analyzed by western blot as described above using primary antibodies against FUS/TLS (1:1000, Proteintech Group, #11570-1-AP), RBM4 (1:1000, Proteintech Group, #11614-1-AP), and HNRPR (1:1000, Proteintech Group, #29980-1-AP).

### Computational Analysis

Conventional molecular dynamics (cMD) simulations were conducted for the following four systems: WT SRC kinase, WT SRC kinase with a phosphorylated Y416 residue, the SRC kinase Y416D variant, and the loop truncated SRC kinase P8E2 variant.

The loop deletions in the P8E2 variant were modelled using AlphaFold2^63^ and Rosetta^78,79^. All systems were simulated in three independent 1 µs replicas, producing 3 µs of analyzed data per system. Simulations were performed with the GPU-accelerated version of GROMACS 2020^80^, using the CHARMM36m force field^81^.

Initial conformations for AlphaFold-seeded MD were generated using the AlphaFold2^63^ multiple sequence alignment (AF2-MSA) subsampling strategy described by Silva et al.^82^, implemented from the public repository. For each variant (WT, P8E2, and selected gatekeeper mutants), 200 AF2 models were generated using default pipeline parameters. Each model was scored with AF2’s predicted local distance difference test (pLDDT) and projected into a reduced conformational space defined by (i) the ΔD coordinate (difference between the distances E310-R409 and K295– E310) and (ii) the activation-loop extension angle (C_α_-atoms of K295–E310–Y/D416). The reduced-space distribution was clustered via k-means (k = 50) in scikit-learn^83^, and one representative conformation from each cluster was selected to seed MD simulations. Selected conformations were simulated for 400 ns each in NAMD using CHARMM36m force field^81^, producing 20 µs of data per system.

Additional computational details on structure preparation and simulation conditions are listed in Supplementary Information. The starting structures as well as analysis and simulation scripts are deposited at the 10.5281/zenodo.17345658.

### Phylogenetic Analysis of the Tyrosine Kinase Domain

A total of 795 protein sequences were collected from UniProt using the enzyme classifications “EC 2.7.10.1” and “EC 2.7.10.2”, covering taxa from Amoebozoa, Filasterea, Choanoflagellata, and Metazoa. For Metazoa, only reviewed sequences were included, while for the other groups, both reviewed and unreviewed sequences were considered. Previously identified fungi-specific tyrosine kinases (239 sequences) were also added to the dataset^84^. All sequences were aligned using MAFFT^85^, and the kinase domains were extracted. CD-HIT was then applied to the isolated kinase domains using a 90% sequence identity cutoff to remove redundancy. The resulting sequences were manually curated to remove those containing large insertions or deletions, followed by realignment using MAFFT. Additionally, 17 kinase domain sequences from human serine/threonine kinases were included as an outgroup. The final dataset, comprising 405 sequences, was used to construct a maximum-likelihood phylogenetic tree using the LG+F+R9 model, which was automatically selected by IQ-TREE^86^.

## Data Availability

The coordinates and structure factors for the P8E2 variant have been deposited in the Protein Data Bank under accession code 9V4E. All raw LC-MS/MS data have been deposited in the ProteomeXchange Consortium via the jPOST repository^66^ under the dataset identifier PXD068232. All computational simulation files, structures and analysis were deposited at 10.5281/zenodo.17345658.

## Code Availability

All custom R scripts used for proteomics data analysis are available upon request.

## Acknowledgments

P.L. gratefully acknowledges funding from the Okinawa Institute of Science and Technology (OIST). P.J. gratefully acknowledges financial support from the JSPS Kakenhi Early Career Scientist Grant (22K15064). S.C.L.K. is the Georgia Research Alliance – Vasser Wooley Chair of Molecular Design at Georgia Tech, and gratefully acknowledges the GRA, Georgia Tech, and the Knut and Alice Wallenberg Foundation (2019.0431) for financial support. We acknowledge the National Academic Infrastructure for Supercomputing in Sweden (NAISS), partially funded by the Swedish Research Council through grant agreement no. 2022-06725, for awarding this project access to the LUMI supercomputer, owned by the EuroHPC Joint Undertaking and hosted by CSC (Finland) and the LUMI consortium. Synchrotron radiation experiments were performed at BL45XU of SPring-8 with the approval of the Japan Synchrotron Radiation Research Institute (JASRI) (proposal no. 2024A2720). We thank John Kuriyan at Vanderbilt University for providing the Peptide Library for High-Throughput Substrate Specificity Screening Assay, and also for the comments and critical reading of the manuscript. We thank Serena Muratcioglu at Vanderbilt University for assisting with the Peptide Library for High-Throughput Substrate Specificity Screening Assay protocol and also with the data analysis. We thank Marco Terenzio at OIST for providing HEK293T cells. We thank Yoshitoshi Hiaro and Fuka Koja from the Instrument Analysis Section (IAS) at OIST for their assistance with proteomics and FACS experiments, respectively. We also thank the IAS and Sequencing Section at OIST for providing instrument access and training. The images of the present work were prepared using Biorender.com.

## Author Contributions

P.L. and P.J. conceived the project and designed the experiments. P.J. performed all the in-vitro and in vivo experiments. P.L. and P.J. analyzed the experimental data. P.J. and B.E.C. solved the X-ray crystal structure. D.Y., A.D., M.R. and S.C.L.K. designed the MD simulations and analyzed the data. R.R.F. and M.O. ran the first round of western blot and analyzed the data. A.I. assisted with or offered guidance regarding proteomics and cell experiments and analysis. P.J. and A.I. analysed the protemics data. P.L. directed and supervised the project. P.L., P.J. and A.I. wrote the manuscript with input from all authors.

---

**Extended Figure 1:**
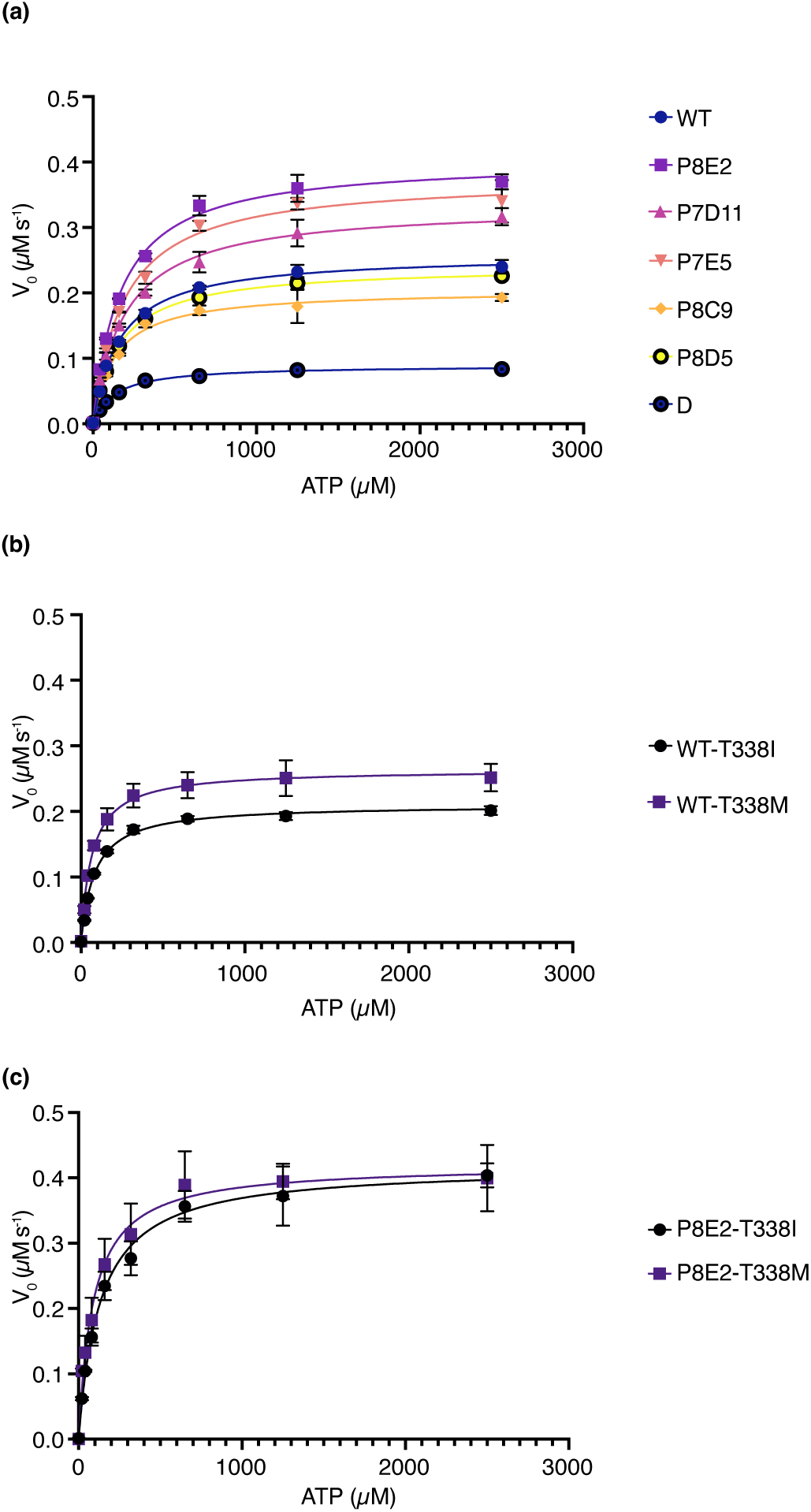
Michaelis-Menten Kinetics of SRC Variants with ATP. Michaelis-Menten plots for SRC and its variants using ATP as the substrate. Initial reaction velocities were measured at varying ATP concentrations for WT, D, and the top five candidates, including P8E2, from high-throughput screening **(a)**, for WT gatekeeper variants **(b)**, and for P8E2 gatekeeper variants **(c)**. Plot represent the mean ± standard deviation (SD) of the initial velocities. Most variants were measured in two independent experiments with duplicates (n = 4), whereas P8E2-T338I and P8E2-T338M were measured in two independent experiments without duplicates (n = 2).

**Extended Figure 2:**
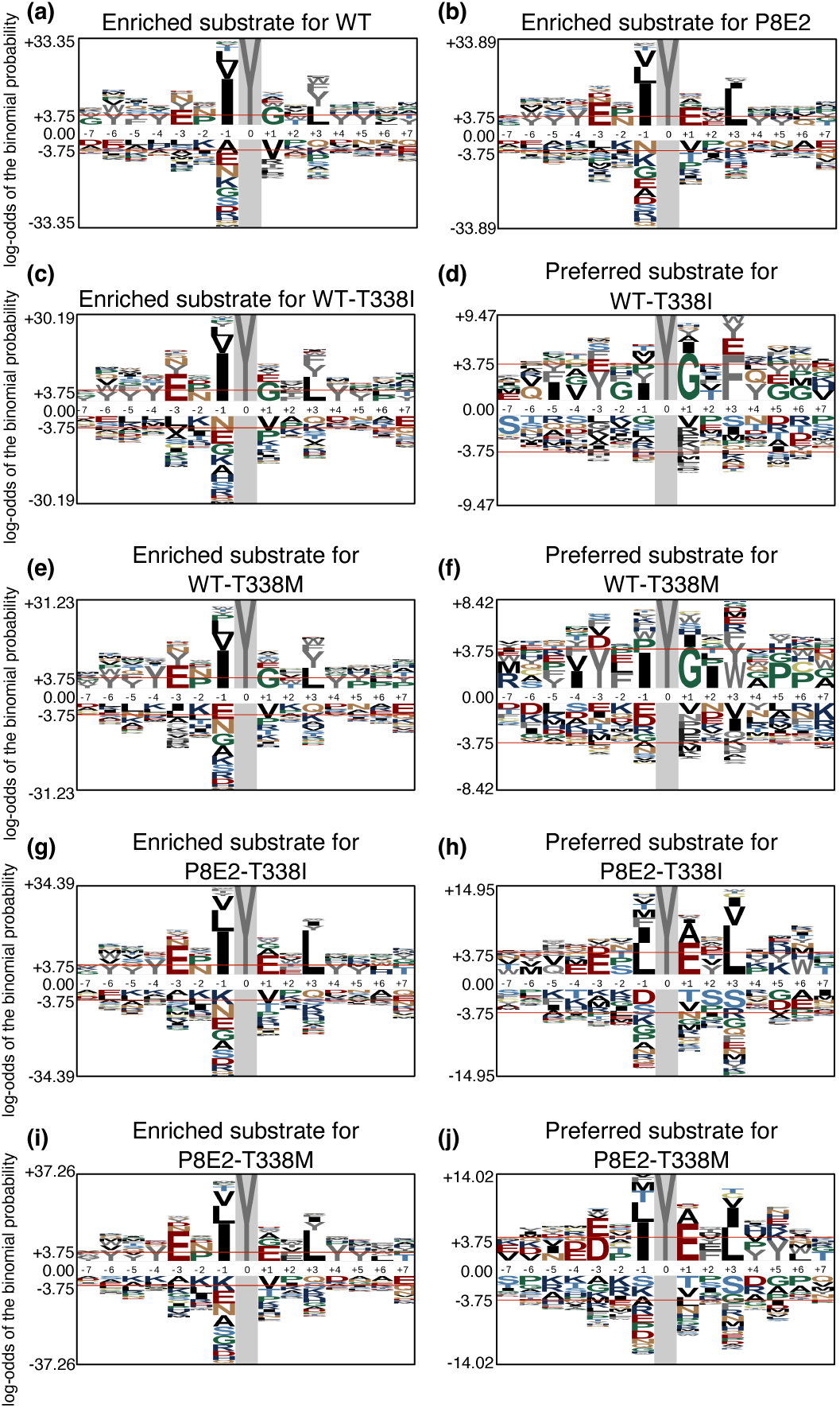
Phospho-Plogo Diagrams Showing Enriched and Preferred Substrate Specificities Across WT, P8E2, and Their Gatekeeper Mutants. Peptides with average enrichment scores >1.5 across three biological replicates were first selected as strong substrates and used to generate the enriched phospho-pLogos. The number of enriched peptides for each variant was: **(a)** WT = 273, **(b)** P8E2 = 260, **(c)** WT-T338I = 248, **(e)** WT-T338M = 298, **(g)** P8E2-T338I = 304, and **(i)** P8E2-T338M= 334. However, enrichment alone does not capture kinase specificity, as many peptides are efficiently phosphorylated by multiple variants. To detect variant-specific preferences, we applied a comparative filtering strategy in which peptides enriched by one variant but not by its counterpart were retained as preferred substrates. For example, for WT-T338I, peptides enriched (>1.5) in P8E2-T338I were removed, and vice versa for P8E2-T338I. The same filtering was applied to the T338M gatekeeper pair and to WT and P8E2, as shown in Figure 1g and 1h. The number of preferred peptides obtained after filtering was: **(d)** WT-T338I = 34, **(f)** WT-T338M = 49, **(h)** P8E2-T338I = 90, and **(j)** P8E2-T338M = 85. Both enriched and preferred phospho-pLogos were generated using the full peptide library (n = 2,587) as background. Residues above the center (0.00 log-odds of the binomial probability) represent overrepresented amino acids, while residues below the center indicate underrepresented residues in the respective enriched or preferred groups.

**Extended Figure 3:**
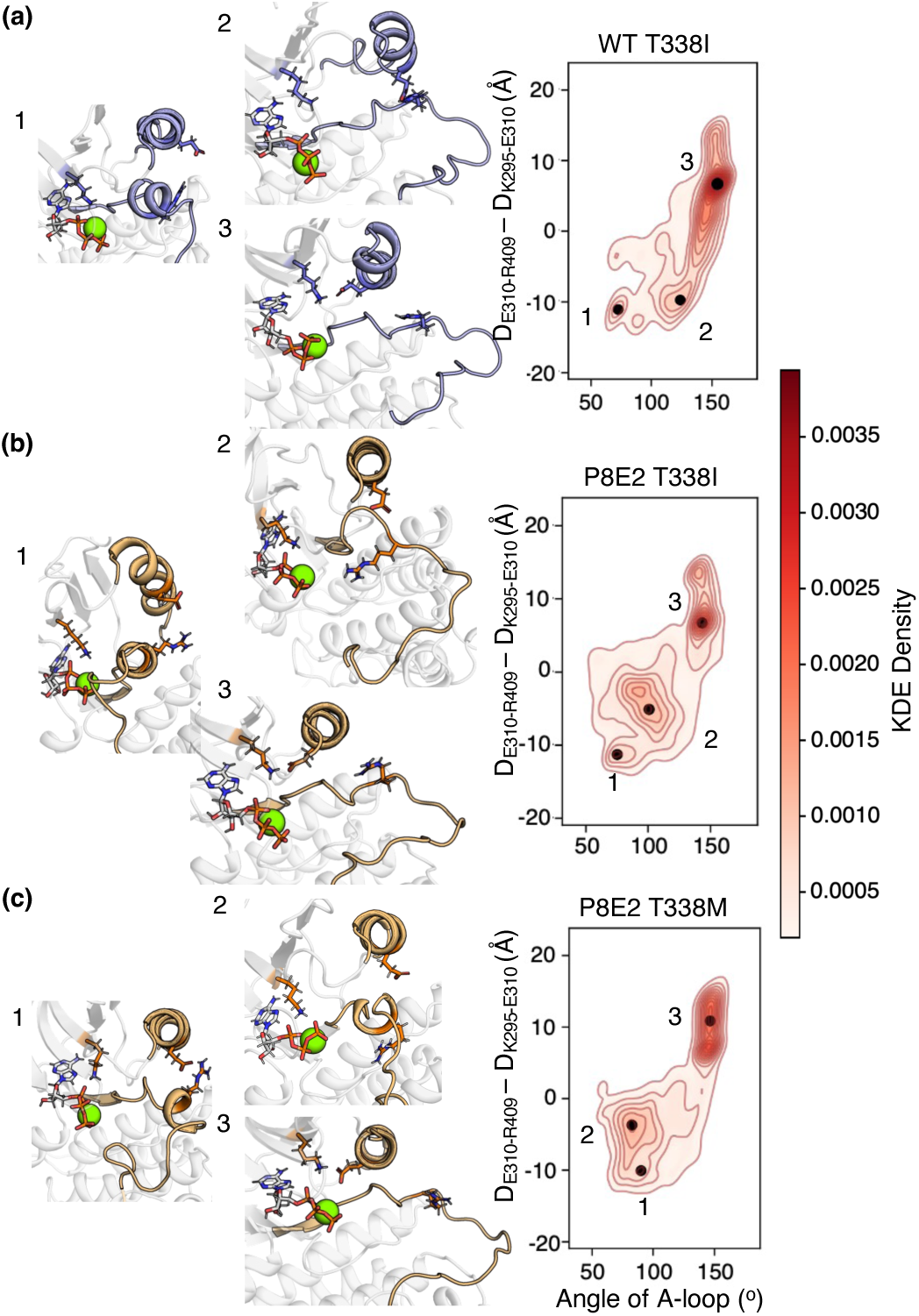
Conformational Landscape of T338 Mutants from AF-Seeded MD Simulations. Kernal density estimate (KDE) plot and selected representative frames of combined data from simulations initialized for AlphaFold2^63^ (AF)-seeded MD for (**a**) WT T338I, (**b**) P8E2 T338I and (**c**) P8E2 T338M mutants. Conformational analysis was based on the difference in distances between E310-R409 and K295-E310, and in the angle between Cα of K295-E310-Y416 for WT-T338I and K295-E310-D416 for P8E2 T338I/M.

**Extended Figure 4:**
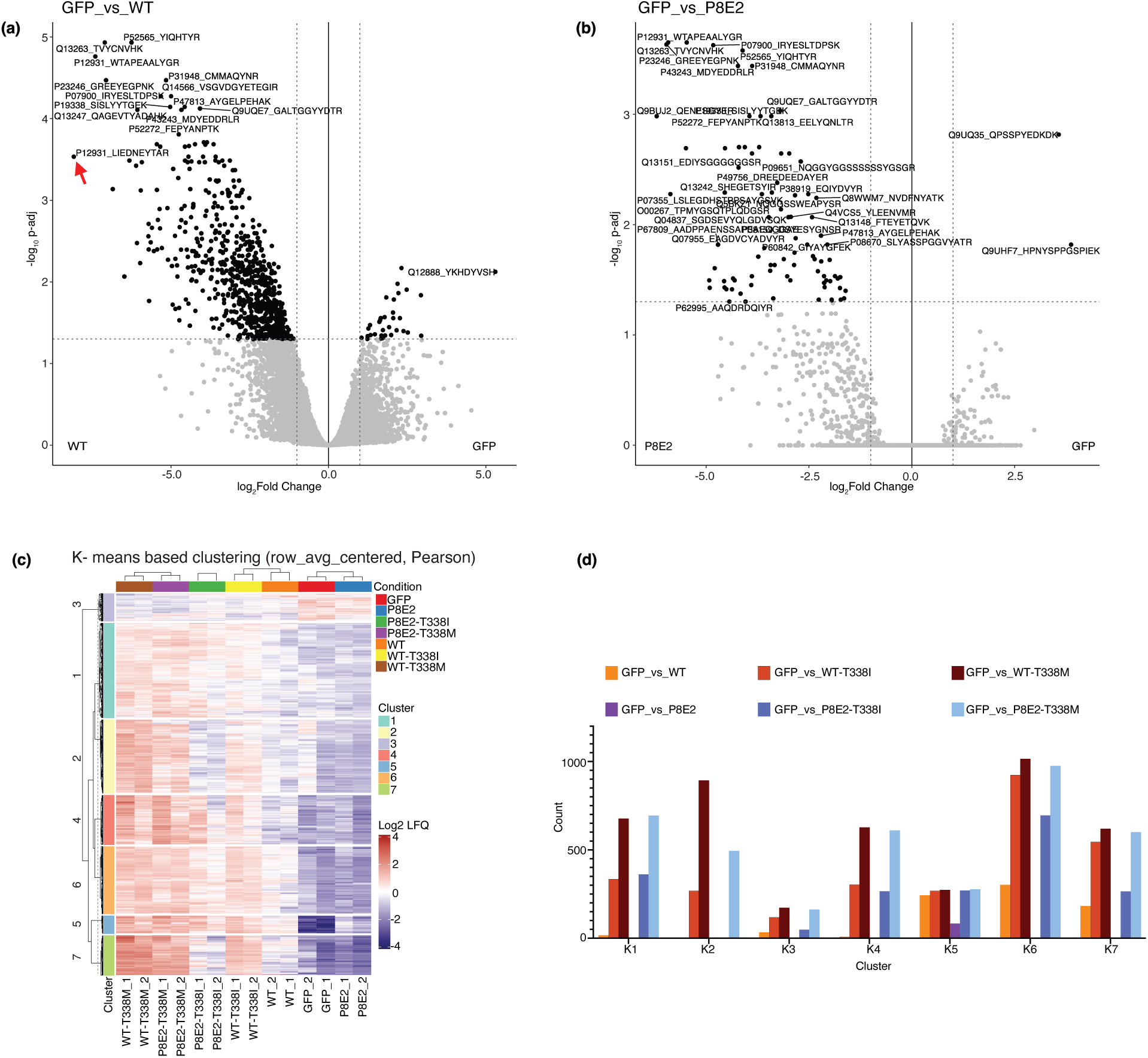
Peptide Level Phosphoproteome Analysis of Differential Expression Across SRC Variants. **(a and b)** Volcano plots show several peptides are differentially expressed. The red arrow indicates that the SRC peptide located in the A-loop is differentially expressed in the GFP_vs_WT SRC **(a)** but not in the GFP_vs_P8E2 contrast set **(b)**, consistent with the fact that the Y416 is substituted in P8E2. **(c)** Hierarchical heatmap for DE peptides with K-means clustering and Peason correlation similar to that of DE proteins (Figure 4c). **(d)** DE peptide counts in each experimental contrast grouped by K-means clusters K1-K7. The trend in this graph is similar to that of Figure 4f (Monte Carlo simulations followed by pairwise chi-squared tests), while the K-means clusters are not aligned to those of the protein clusters due to the stochastic nature of K-means clustering. For example, here K5 is similar to K4 for protein clusters in Figure 4f.

**Extended Figure 5:**
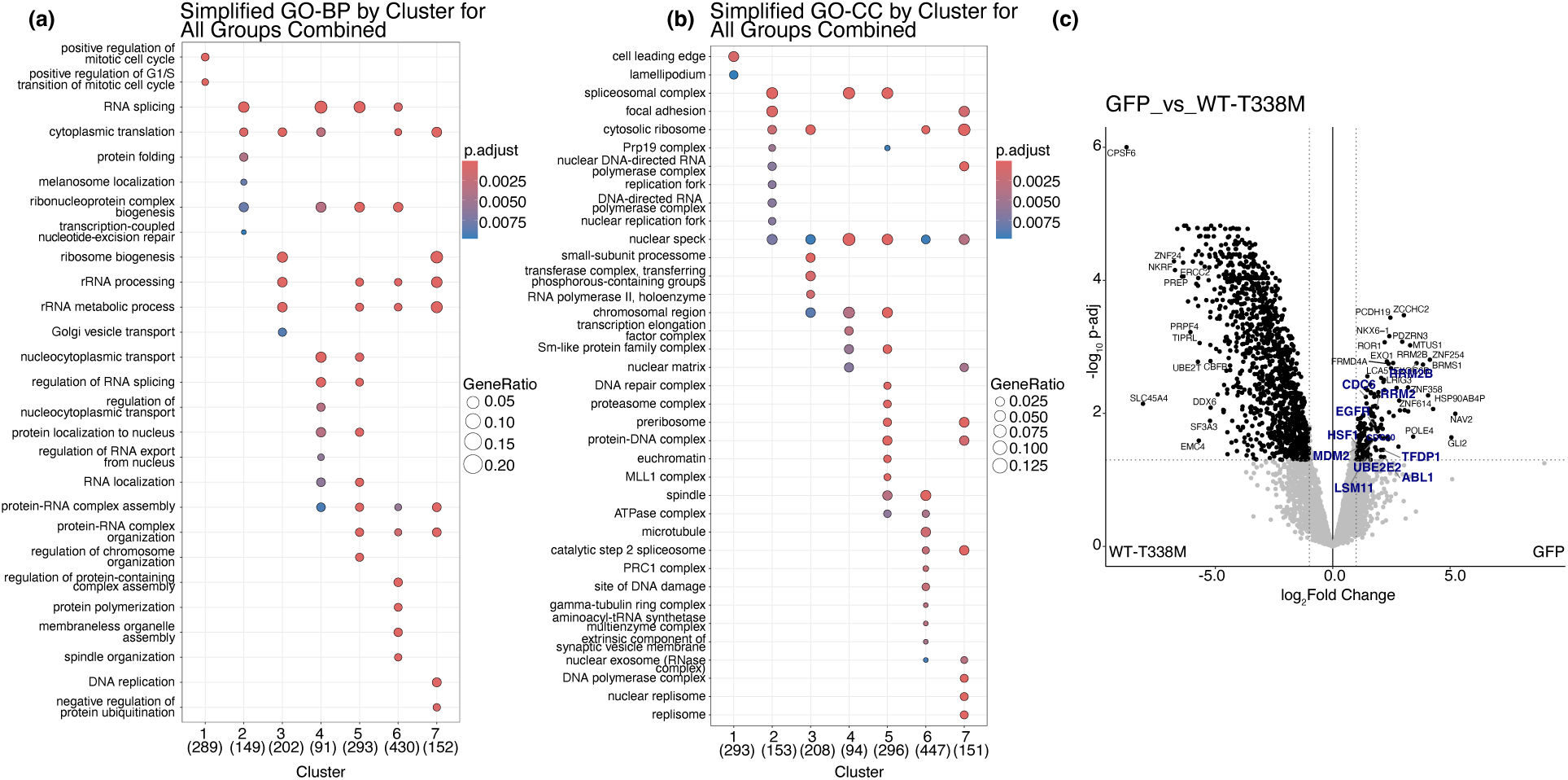
Functional Enrichment of K-Means Clusters Reveals Signaling Rewiring by SRC Variants. **(a and b)** Enrichment analysis is performed using Gene Ontology Biological Process (GO-BP) **(a)** and Cellular Component (GO-CC) **(b)** databases for seven K-means clusters without separating the experimental contrast sets. These panels therefore provide functional identities to K-means clusters shown in Figures 4c (proteins) and Extended Figure 4c. K-means clusters for proteins and peptides do not align with each other due to the stochastic nature of K-means clustering. **(c)** Volcano plot is generated for the contrast set of GFP vs WT-T338M to highlight the proteins associated with the GO-BP term of ‘positive regulation of mitotic cell cycle’ in K1 cluster (labels with bold blue font), including EGFR, as downregulated by WT-T338M (i.e., greater in the GFP control).

**Extended Figure 6:**
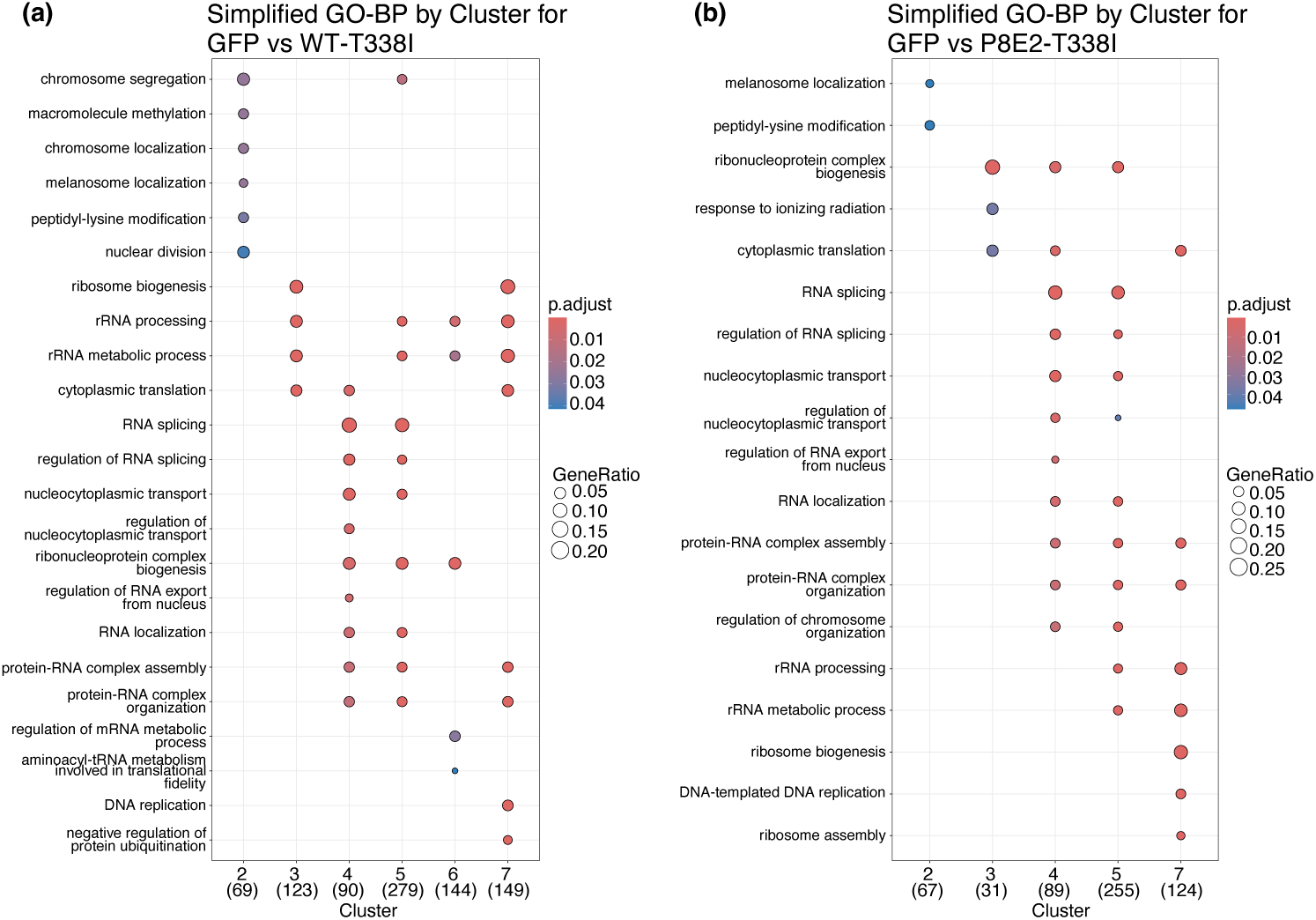
GO-BP enrichment of protein clusters for T338I mutants. **(a and b)** Enrichment analysis is performed using GO-BP database for seven K-means clusters after separating DE proteins in the contrast sets GFP vs WT-T338I **(a)**, GFP vs P8E2-T338I **(b)**. These enrichment patterns align with those in Figure 4f and indicate a lower overall induction of DE proteins by the T338I variant compared to its T338M counterpart.

**Extended Figure 7:**
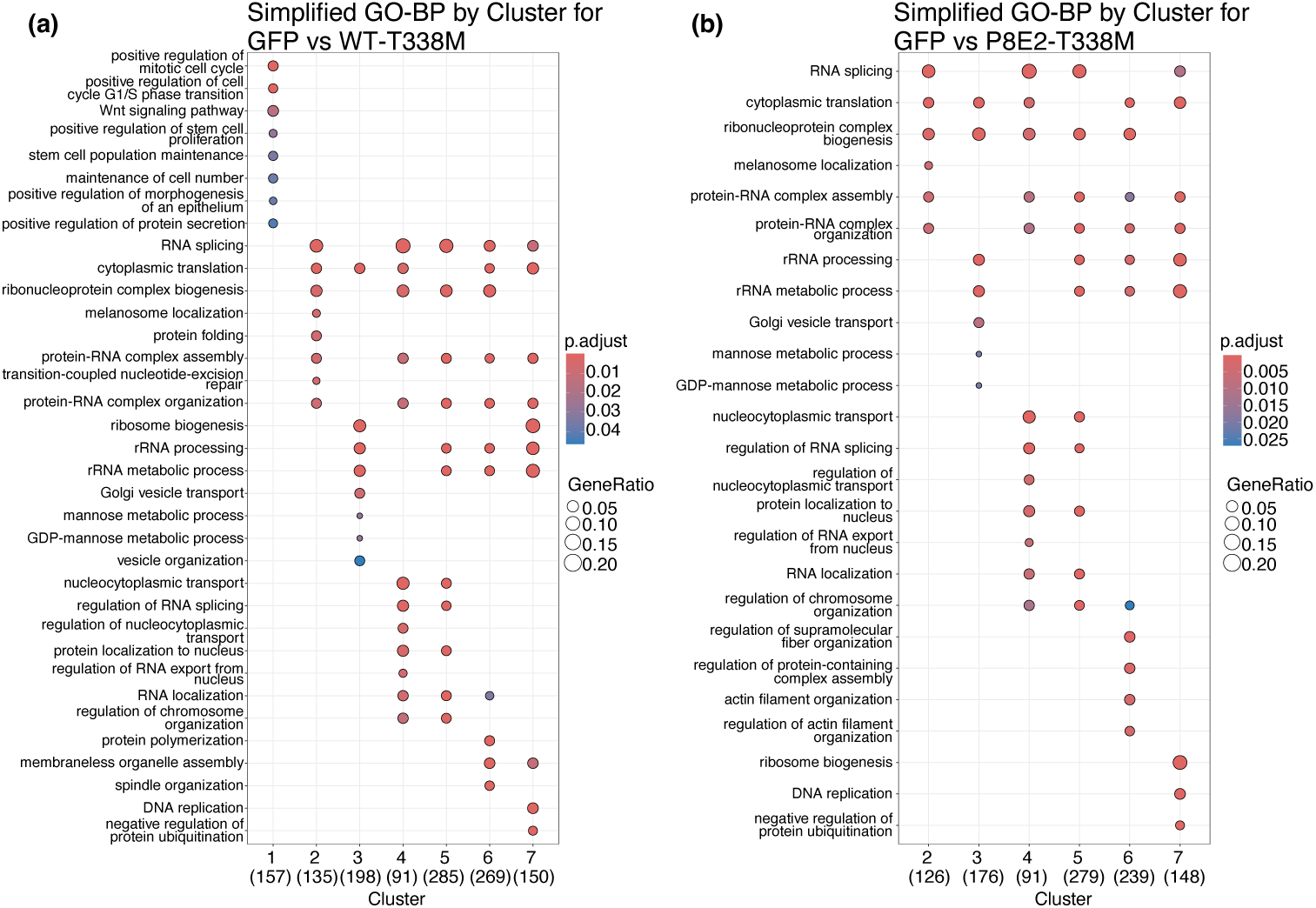
GO-BP enrichment of protein clusters for T338M mutants. **(a and b)** Enrichment analysis is performed using GO-BP database for seven K-means clusters after separating DE proteins in the contrast sets GFP vs WT-T338M **(a)**, GFP vs P8E2-T338M **(b**). Consistent with Figure 4f, the T338M gatekeeper mutation shows a broader induction of DE proteins than the T338I variant in both the WT and P8E2 backgrounds.

## Notes

### Competing Interest Statement

The authors have declared no competing interest.

