## Supplementary material for "Minimal Perturbation of Activation Loop Dynamics Rewires Kinase Signaling": Table S1

### Supplementary Text

#### Supplementary Note 1: Evolutionary Context of Autophosphorylation Sites

The Y416D substitution described in the main text was designed to test how acidic residues can mimic phosphorylation at key regulatory sites. To provide context for this design, comparative genomic analyses suggest that some conserved autophosphorylation sites across the kinome may have originated from ancestral acidic residues, such as aspartate or glutamate, that were later replaced by phosphorylatable residues. This transition is thought to have enabled tighter and more dynamic regulatory control of kinase activity. Ferrell and colleagues estimated that roughly 5% of phosphosites may have evolved from acidic residues, both during the divergence of eukaryotes from prokaryotes and in subsequent evolutionary events[88]. While this trend was most evident for serine sites, likely reflecting their overall abundance[89], it does not exclude the possibility that tyrosine autophosphorylation sites followed a similar trajectory. Although relatively few studies have examined the evolution of phosphotyrosine sites specifically, their emergence may have been crucial for supporting the increasingly complex signaling functions required in multicellular organisms[90].

#### Supplementary Note 2: Structural and Functional Effects of Gatekeeper Mutations in SRC

The conserved gatekeeper residue T338 lies within the ATP-binding site and has been a major focus in kinase biology and drug design[91,92]. Mutations at this site, particularly T338I and T338M, are frequently observed in cancers and confer resistance to ATP-competitive kinase inhibitors[93,94]. Mechanistically, these substitutions reinforce the hydrophobic regulatory spine and destabilize the inactive conformation, thereby shifting the equilibrium toward the active state and promoting both autophosphorylation and catalytic activity.

Although the gatekeeper residue is spatially distant from the activation loop and does not directly contact substrates, its mutation induces coordinated structural changes in both the regulatory spine and the activation loop. This expands the conformational landscape of the kinase and can influence substrate selection. We hypothesized that this enhanced flexibility might synergize with activation loop alterations to refine specificity.

However, contrary to this hypothesis, gatekeeper mutations in the P8E2 background obscured the substrate preference shifts observed in the parent variant under cellular conditions. In vitro assays revealed similar substrate profiles to P8E2 but with reduced exclusivity. While the enhanced activity of the WT gatekeeper mutant can be explained by increased autophosphorylation, the similar effect in the deletion variant suggests that these mutations also reshape activation loop conformation. Conformational landscape analysis further supported this view: both P8E2-T338I and P8E2-T338M variants converged on catalytically competent states with extended loop conformations.

Together, these findings indicate that while activation loop sequence is a primary determinant of substrate specificity, fine-tuning the broader conformational dynamics of the kinase is essential to fully realize this specificity.

**Supplementary Computational Methodology Details**

A two-fold computational approach was used to characterize the effects of residue deletions in the A-loop and gatekeeper mutations of the SRC kinase variants. First, conventional molecular dynamics (MD) simulations were utilized to assess the localized effects of the relevant amino acid substitutions on the flexibility of the A-loop. Further, our MD simulation approach was combined with the AlphaFold2[95] sequence clustering algorithm[96] to generate additional starting conformations and compare coordinated dynamics between the A-loop and αC-helix among the variants.

<u>*Conventional Molecular Dynamics*</u>

Conventional molecular dynamics (cMD) simulations were conducted for the following four systems: wild-type (WT) SRC kinase, WT SRC kinase with a phosphorylated Y416 residue, the SRC kinase Y416D variant, and the loop truncated SRC kinase P8E2 variant.

The loop deletions in the P8E2 variant were modelled using AlphaFold2[95] and Rosetta[97,98]. 10 initial structures of the deletion mutants were generated using AlphaFold2 Monomer. Templates were sourced from the UniRef90[99], MGnify[100], Uniclust30[101], PDB70[102], and BFD[95] databases. AlphaFold2[95] was used to perform an initial relaxation of these structures. Following this, RosettaRelax[103] was used to locally relax and rank the mutant structures, and the lowest ranked structure was selected for subsequent work.

All systems were simulated in three independent 1 μs replicas, producing 3 μs of analyzed data per system. Simulations were performed with the GPU-accelerated version of GROMACS 2020[104], using the CHARMM36m[105] force field along with the TIP3P[106] water model. System preparation was performed using the standard protocol of the CHARMM-GUI[107] builder, with the phosphorylated pY416 parameterized using CHARMM-GUI[107] solution builder tools. Each system was solvated in a cubic box of 67x67x67 Å or 10 Å from the edge boundary of the protein with 0.15 M NaCl added to the system.

Each system underwent energy minimization using the steepest descent algorithm, followed by equilibration in the NVT ensemble. Temperature was maintained at 303.15 K using the Nosé-Hoover[108,109] thermostat. Production MD was run using the Nosé-Hoover thermostat[108,109] and pressure was regulated at 1 atm using the Parrinello-Rahman barostat[110] with isotropic coupling and a relaxation time of 5.0 ps. All bonds involving hydrogen atoms were constrained using the LINCS[111] algorithm, enabling a 2 fs integration time step. Electrostatics were treated using the Particle Mesh Ewald (PME) method[112], with a Verlet cutoff scheme[113] and a 1.2 nm cutoff for both Coulomb and van der Waals interactions.

Root mean square fluctuation (RMSF) values were computed for the $C_\alpha$-atoms of each residue with reference to the initial structure for each simulation set using MDAnalysis 2.7.0[114,115]. To show the flexibility of the A-loop, RMSF values for this region were projected onto the first 50 structural clusters, derived from clustering analysis performed using the single linkage clustering protocol with a cutoff of 0.8 Å, as implemented into GROMACS[116] 2020. Clustering was performed on the concatenated trajectories across all three replicas, considering all protein atoms in the structural comparison. The resulting representative structures were visualized in PyMOL Molecular Graphics System, Version 2.6.2 Schrödinger, LLC., with the A-loop colored according to per-residue RMSF values.

The $\chi_1$ and $\chi_2$ dihedral angles of residue F424 (A-loop, P+1 loop region) were computed for each frame of the conventional MD trajectories using MDAnalysis 2.7.0[114,115]. Two-dimensional histograms were generated with $100 \times 100$ bin resolution. Rotamer states were classified according to standard $\chi_1$ ranges: gauche- ($300° \pm 30°$), gauche+ ($60° \pm 30°$), and trans ($180° \pm 30°$). The population of each rotamer was computed as the fraction of simulation frames falling into the corresponding basin.

*AlphaFold2-Initialized Molecular Dynamics*

Initial conformations for enhanced sampling were generated using the AlphaFold2[95] multiple sequence alignment (AF2-MSA) subsampling strategy described by Silva et al.[117], implemented from the public repository. For each variant (WT, P8E2, and selected gatekeeper mutants), 200 AF2 models were generated using default pipeline parameters. Each model was scored with AF2's predicted local distance difference test (pLDDT) and projected into a reduced conformational space defined by (i) the ΔD coordinate (difference between the distances E310-R409 and K295–E310) and (ii) the activation-loop extension angle ($C_\alpha$-atoms of K295–E310–Y/D416). The reduced-space distribution was clustered via K-means ($k = 50$) in scikit-learn[118], and one representative conformation from each cluster was selected to seed MD simulations.

System preparation for each AF2-derived starting structure was performed in VMD 1.9.4a57 using the psfgen plugin. The CHARMM36m force field[119] was used for the protein, with ATP parameters obtained from the CHARMM-GUI[107] Ligand & Solution Builder. Systems were solvated in a rectangular TIP3P[106] water box, with at least 12 Å padding from any solute atom, neutralized, and ionized to 0.15 M NaCl concentration. Energy minimization (5000 steepest descent steps) was followed by NVT equilibration with backbone restraints (250 kcal $mol^{-1}$ Å$^{-2}$), gradually released over a series of 10 ps intervals ($250 \rightarrow 25 \rightarrow 2.5 \rightarrow 0.25$ kcal $mol^{-1}$ Å$^{-2}$). An additional 1 ns unrestrained NPT equilibration was performed prior to production. All simulations were run in NAMD with a 12 Å non-bonded cutoff, switching at 10 Å, PME electrostatics[112], and a 2 fs integration timestep. Temperature was maintained at 298 K via Langevin dynamics (damping constant = 1 $ps^{-1}$), and pressure at 1 atm with a Langevin barostat[120] (period = 100 fs, decay = 50 fs). Each selected starting structure was simulated to produce a 400 ns trajectory.

From each AF2-seeded MD trajectory, conformational space was described in terms of two features: (i) the difference between the distances E310-R409 and K295-E310, and (ii) the activation-loop extension angle K295-E310-Y/D416. The distance measurement was based on the closest terminal sidechain oxygen and nitrogen atoms.

### Supplementary Figures

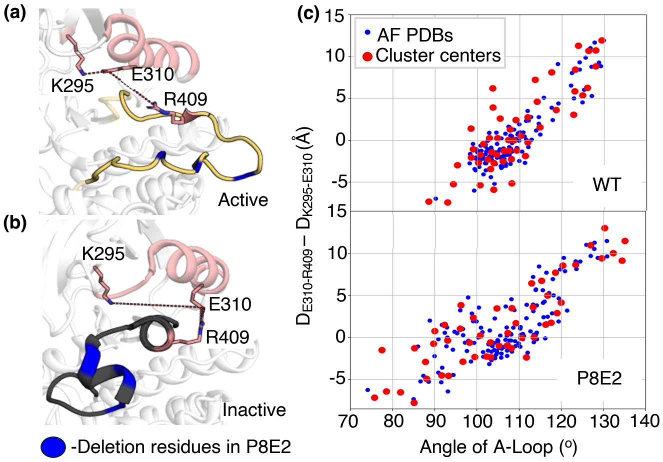

**Figure S1**: **Structural Determinants Distinguishing Active and Inactive SRC Kinase Conformations. (a and b)** Reference crystal structures that define the two variables used to distinguish active and inactive conformation of SRC kinase. (**a**) In the active state (PDB ID: 3DQW[121]) the *C*-helix (pink) is pulled inward, allowing the catalytic E310–K295 salt bridge (red dashed line) to form while R409 disengages from E310; permitting the activation loop (yellow) to fully extend. (**b**) In the inactive kinase (PDB ID: 3U4W[122]), R409 caps E310, the E310–K295 contact is broken, and the A-loop folds back over the active site. Residues deleted in P8E2 (N414, T417, R419) are highlighted in blue. (**c**) Distribution of 200 AlphaFold2[95] models (blue) and their K-means cluster centers (red) projected on the two coordinates: A-loop angle between K295-E310-Y/D416 and the distance difference $\Delta D = D_{E310\text{-}R409} - D_{K295\text{-}E310}$. Positive ΔD marks active-like conformations (E310–K295 short, R409–E310 long); negative values mark inactive-like geometries.

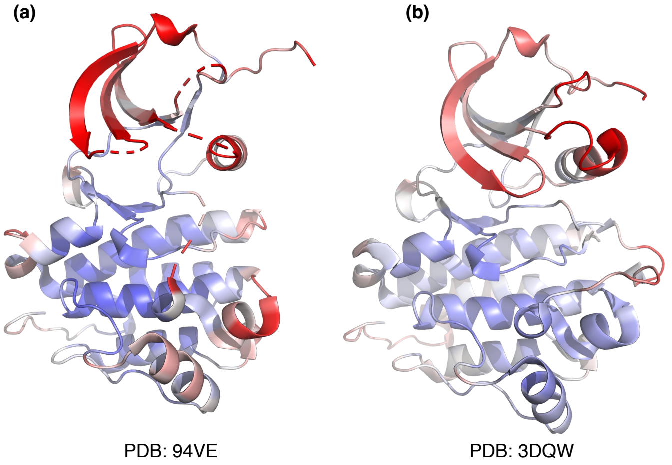

**Figure S2: Structural Comparison of P8E2 and WT SRC Kinase Colored by B-Factor.** Cartoon representation of **(a)** P8E2 (PDB 9V4E; this work) and **(b)** WT-T338I SRC kinase in the active conformation (PDB 3DQW[121]), colored according to B-factor (Å²). Low B-factors are shown in blue, intermediate B-factors in white, and high B-factors in red (range 10–50 Å²). P8E2 exhibits elevated B-factors at the N- and C-termini, and the activation loop is unresolved, highlighting its dynamic and flexible nature compared to the ordered active conformation of WT SRC kinase.

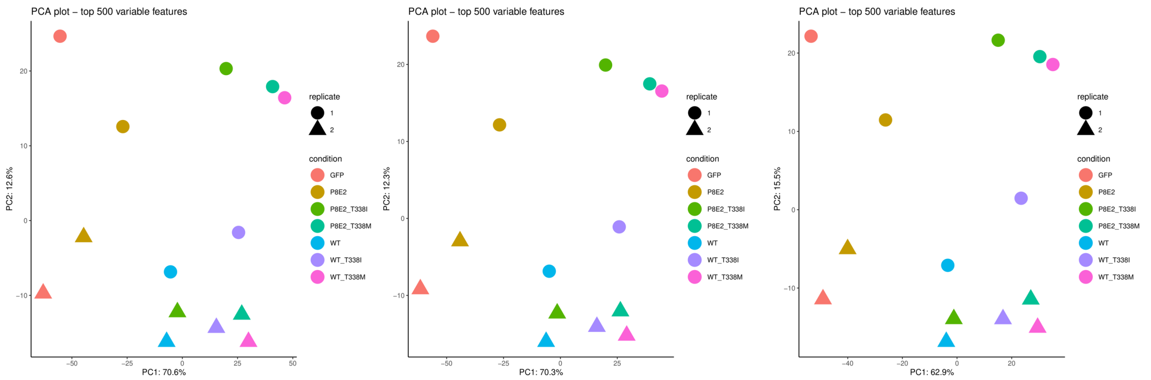

bpca_(no filter)_pro bpca_(prefilter50)_pro bpca_(vsn_prefilter50)_pro

**Figure S3: Principal Component Analysis (PCA) of Proteomics Data with Different Imputation Strategies**. PCA was performed to check the proteome data and to compare the data processing workflow of 'bpca_(no filter)_pro', 'bpca_prefilter50_pro', or 'bpca_vsn_prefilter50_pro' indicated in Table S2. PC2 scaled mainly between the two replicates of each experimental group, while PC1 spread over the experimental groups. Choices of processing workflow did not visibly affect PCA.

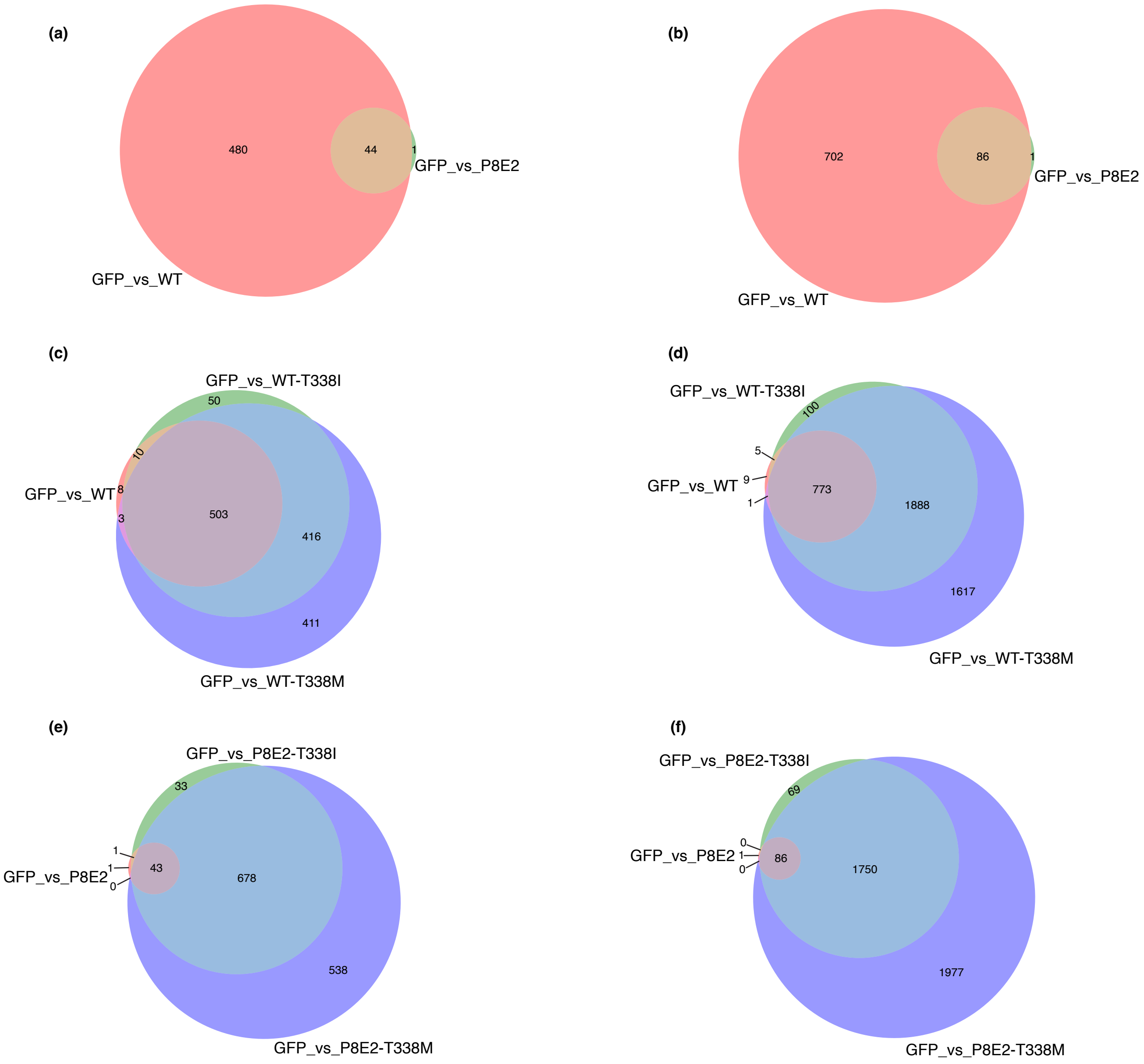

**Figure S4**: **Venn Diagrams of Differentially Expressed Proteins and Peptides Across Kinase Variants**. Left panel **(a,c,e)** DE proteins; Right panel **(b,d,f)**: DE peptides. **(a and b)** Comparison of WT and P8E2. **(c and d)** Comparison of WT, WT-T338I, and WT-T338M. **(e and f)** Comparison of P8E2, P8E2-T338I, and P8E2-T338M.

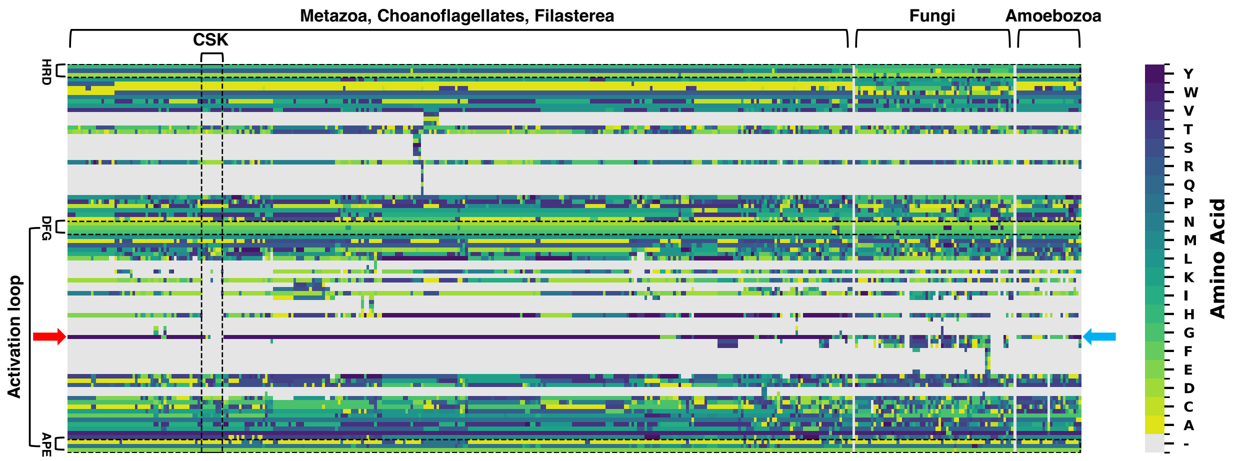

**Figure S5: Multiple Sequence Alignment of Eukaryotic Kinase Domains**. Heatmap of the multiple sequence alignment focused on the activation loop region. Sequences are arranged horizontally. A conserved motif is observed across all sequences. Canonical autophosphorylation sites are marked with red and blue arrows. CSK sequences, characterized by shorter activation loops, are outlined with black dotted lines.

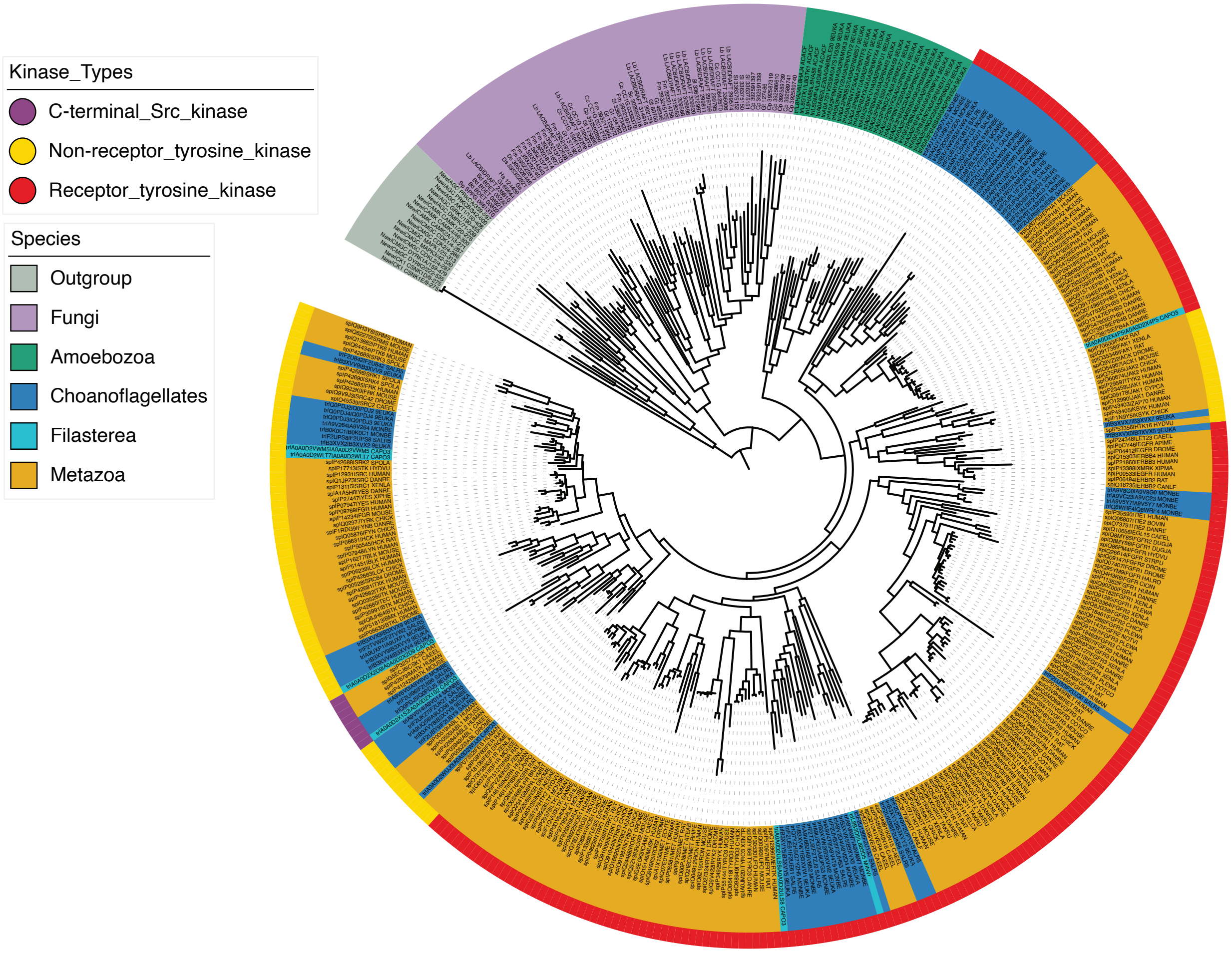

**Figure S6: Phylogenetic Tree of Tyrosine Kinase Domains Across Eukaryotic Lineages.** Phylogenetic tree of the kinase domain across diverse species. Metazoa, Choanoflagellates, and Filasterea form distinct clades that correspond to Receptor Tyrosine Kinases, Non-Receptor Tyrosine Kinases, and C-terminal Src Kinase (CSK). Tyrosine kinase subtypes are not clearly distinguishable in Amoebozoa and Fungi, where no canonical TK clades emerge, although amoebozoan kinases cluster relatively close to those of premetazoa and metazoan.

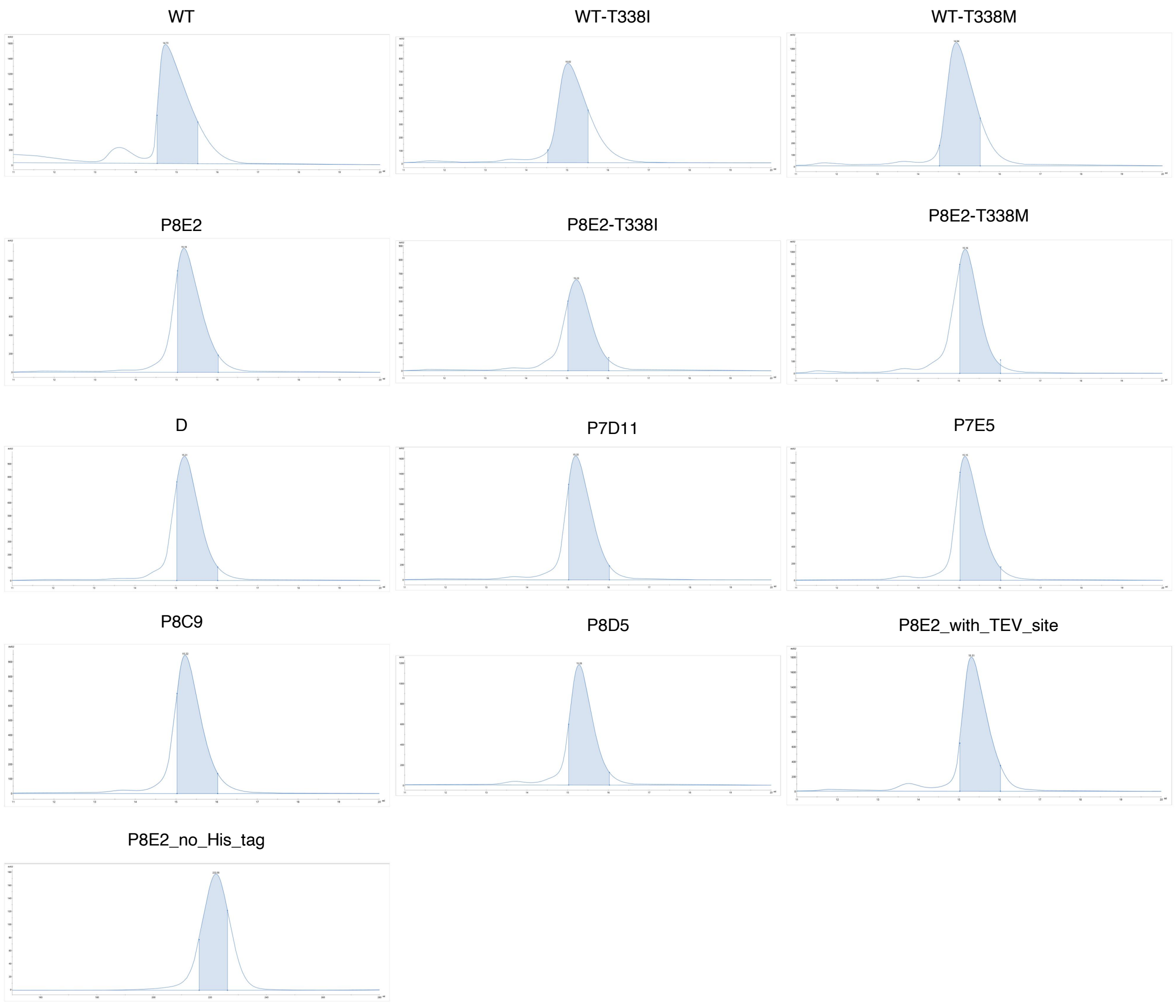

**Figure S7**: **Size Exclusion Chromatography (SEC) Profiles of SRC Variants**. SEC profiles of SRC variants obtained using a Superdex 200 10/300 column. The elution range for all variants is between 14–16 mL. The P8E2 variant with the His-tag cleaved was run on a HiLoad 26/600 Superdex 75 pg column and eluted between 210–230 mL. Shaded regions indicate the fractions that were analyzed by SDS-PAGE.

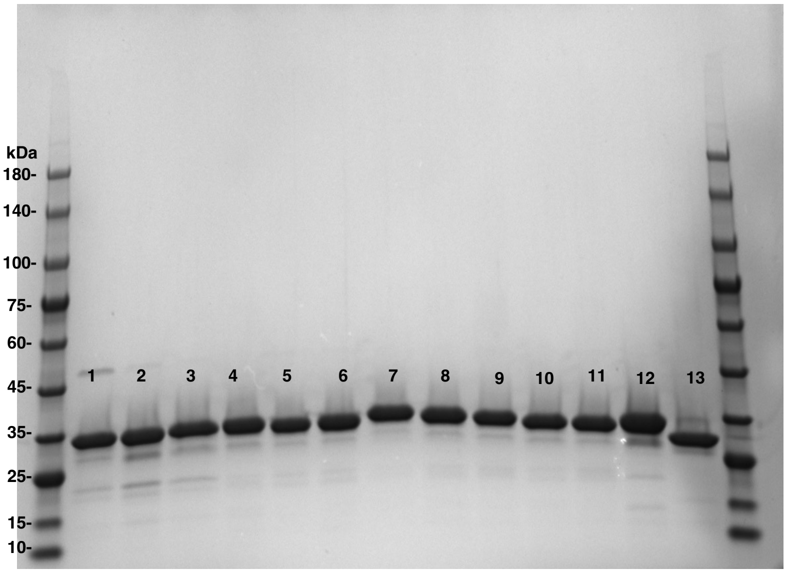

Lane:
1. WT
2. WT-T338I
3. WT-T338M
4. P8E2
5. P8E2-T338I
6. P8E2:T338M
7. D
8. P7D11
9. P7E5
10. P8C9
11. P8D5
12. P8E2_with_TEV_site
13. P8E2_no_His_tag

**Figure S8: SDS-PAGE Analysis of SRC Kinase Variants**. Approximately 5 µg of each purified protein was loaded per well and electrophoresed at 220 V for 30 minutes to assess protein purity and relative molecular weight.

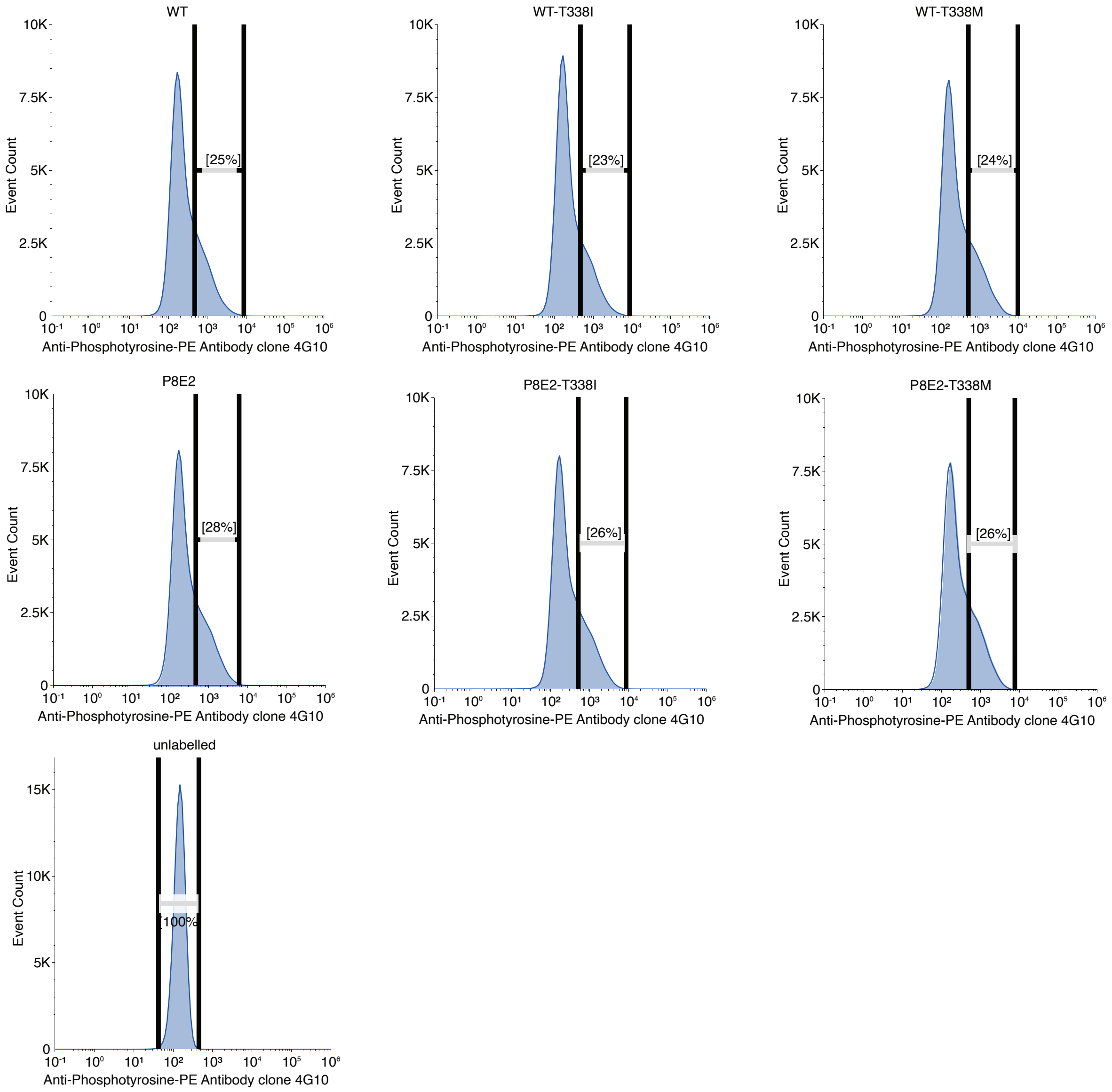

**Figure S9. Flow Cytometry Histograms of WT SRC, P8E2, and Gatekeeper Variants.** Representative histogram (1 of 3 biological replicates) for WT SRC, P8E2, and gatekeeper variants. Approximately 100,000 events were recorded to generate the histogram, and 2–3 million cells were collected from a region offset by 25% from the gated population for next-generation sequencing. Raw flow cytometry data were processed and plots were generated using Floreada.io (https://floreada.io). The X-axis indicates fluorescence detected using the anti-phosphotyrosine antibody 4G10 (PE-conjugated), and the Y-axis shows event count per fluorescence bin.

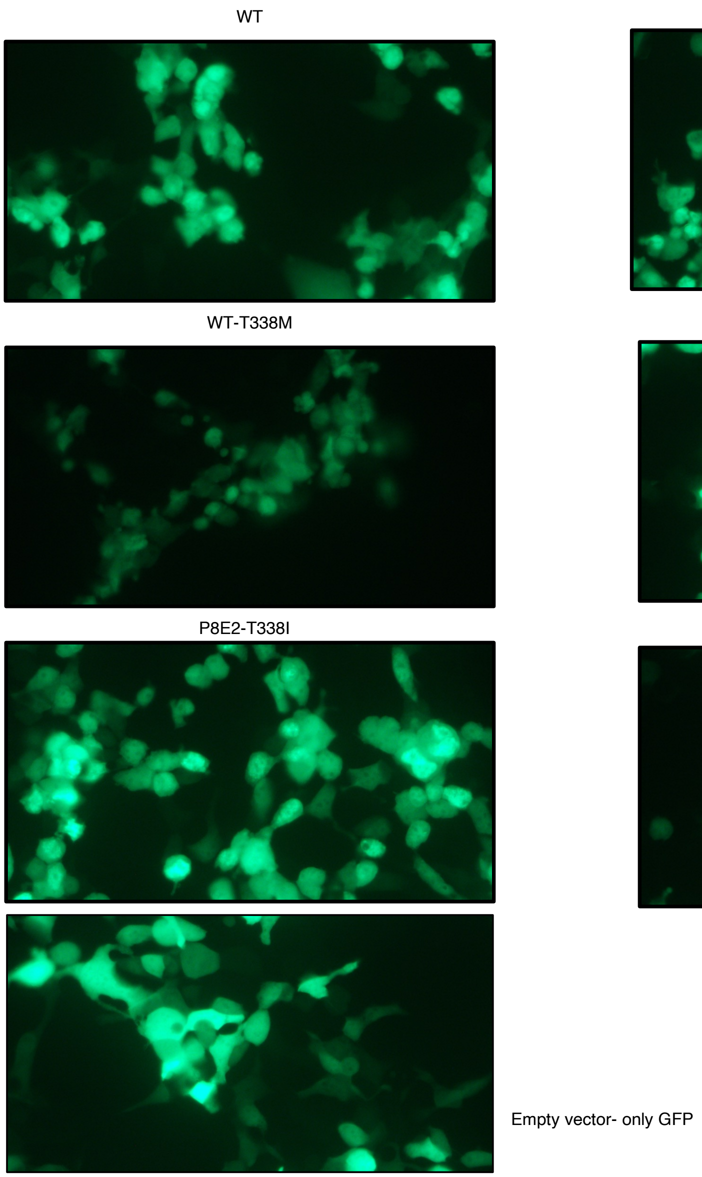

**Figure S10: Fluorescence Images ff HEK293T Cells Expressing GFP-Tagged WT SRC, P8E2, and Gatekeeper Mutants**. Representative Fluorescence images (1 of 2 biological replicates) of HEK293T cells transiently expressing WT SRC, P8E2, and their respective gatekeeper mutants with N-terminal GFP tags. Images were captured at 40× magnification and correspond to the samples used in the proteomics experiment.

### Supplementary Tables

**Table S1: Activation Loop Variants of SRC Kinase.** Positions in the activation loop where residues were either substituted or deleted. Each of the six WT residues (L410, N414, T417, A418, R419, Q420) was subjected to one or more amino acid substitutions or complete deletion (Δ) in various combinations to evaluate their effects on catalytic activity. A full list of all designed variants is provided in the Supplementary Data 2

| Position (WT) | Substituted Residue(s) |
|---|---|
| L410 | V, I, M, Δ |
| N414 | D, S, T, Δ |
| T417 | N, L, V, M, E, S, Δ |
| A418 | P, Δ |
| R419 | Q, H, Δ |
| Q420 | E, V, Δ |

**Table S2: Effects of Prefilter, Normalization, and Missing Value Imputation (MVI) on the Number of DE Peptides and Proteins.** A custom filter, 'prefilter50', removes the peptide if NA is found as a missing value in more than 50% of the 14 samples (7 groups, each with 2 replicates). The prefilter removes approximately by 39% or 17% of total peptides or proteins (the suffix 'pep' or 'pro'), respectively . *Vsn* is "*variance stabilization normalization*"[123,124]. *Bpca* is "*Bayesian PCA missing value estimation*"[125]. *FragPipeAnalystR* evaluates differential expression with *limma* "*Linear Models for Microarray and Omics Data"*[126]. Significance is based on FDR/BH (*p_adj*) < 0.05 and | log2FC | > 1.0. The percentage in parentheses are percent total. The processing name was given in the chronological order that a filter or normalization was applied. For our analysis of DE peptides or proteins, we chose '*bpca_vsn_prefilter50*' in the workflow unless otherwise indicated.

| Processing Name | Significant Peptides or Proteins | Total |
|---|---|---|
| bpca_(no filter)_pep | 8,687 (28%) | 31,105 |
| bpca_(no filter)_pro | 2,295 (36%) | 6,334 |
| bpca_prefilter50_pep | 5,811 (31%) | 18,916 |
| bpca_prefilter50_pro | 1,860 (35%) | 5,272 |
| vsn_bpca_prefilter50_pep | 5,640 (30%) | 18,916 |
| vsn_bpca_prefilter50_pro | 1,837 (35%) | 5,272 |
| bpca_vsn_prefilter50_pep | 5,787 (31%) | 18,916 |
| bpca_vsn_prefilter50_pro | 1,682 (32%) | 5,272 |

**Table S3: Phosphopeptides Identified in SRC Family Proteins.** *The letter Y in bold typeface indicates the phosphorylated tyrosine residue in the peptide sequence. The red Y indicates the autophosphorylated tyrosine residue in the A-loop. The peptide sequence is not unique to the protein identified in the proteomics software FragPipe as NCBI BLASTP identifies other SRC members share the peptide sequence.

| Peptide Sequence* | IDed by FragPipe | BLASTP ClusteredNR_100% | Significant in comparisons of GFP vs: |
|---|---|---|---|
| LIEDNE**Y**TAR | SRC | FYN, YES, LCK | WT, T338I, T338M |
| VP**Y**PGMVNR | SRC | YES | P8E2-T338I, P8E2-T338M, WT, WT-T338I, WT-T338M |
| WTAPEAAL**Y**GR | SRC | FYN, YES | P8E2, P8E2-T338I, P8E2-T338M, WT, WT-T338I, WT-T338M |
| VIEDNE**Y**TAR | LYN | HCK | P8E2-T338I, P8E2-T338M, WT-T338I, WT-T338M |

**Table S4: X-Ray Data Collection and Refinement Statistics**

| **Structure** | P8E2 |
|---|---|
| PDB code | 9V4E |
| **Data collection** | |
| Space group | $P2_12_12_1$ |
| Cell dimensions | |
| *a*, *b*, *c* (Å) | 41.5, 62.9, 105.4 |
| α, β, γ (°) | 90.0, 90.0, 90.0 |
| Resolution (Å) | 41.46 – 1.78 (1.89 – 1.78)* |
| $R_{merge}$ (%) | 8.4 (73.1) |
| $CC_{1/2}$ (%) | 99.9 (85.2) |
| $I / \sigma I$ | 19.52 (2.79) |
| Completeness (%) | 99.8 (99.3) |
| Redundancy | 8.1 (8.4) |
| **Refinement** | |
| Resolution (Å) | 40.87 – 1.78 |
| No. unique reflections | 27179 |
| $R_{work}$ / $R_{free}$ | |
| No. atoms | |
| Protein | 1925 |
| Ligand/ion | 32 |
| Water | 94 |
| *B*-factors ($Å^2$) | |
| Protein | 32.7 |
| Ligand/ion | 61.7 |
| Water | 34.0 |
| R.m.s. deviations | |
| Bond lengths (Å) | 0.009 |
| Bond angles (°) | 1.72 |

*Values in parentheses are for highest resolution shell.
